# Children and adults use distinct neurocognitive mechanisms to support successful memory-based inference

**DOI:** 10.64898/2026.04.21.719709

**Authors:** Christine Coughlin, Margaret L. Schlichting, Neal W Morton, Katherine R. Sherrill, Michelle Moreau, Alison R. Preston

**Affiliations:** Department of Psychology, University of Texas at Austin, 108 East Dean Keeton Street, Austin, TX 78712; Department of Psychology, University of Toronto, 100 St. George Street, Toronto, ON M5S 3G3; Department of Neuroscience, University of Texas at Austin, 1 University Station, Stop C7000, Austin, TX 78712

**Keywords:** inference, hippocampus, posterior parietal cortex, neural geometry, middle childhood, early adolescence

## Abstract

Reasoning depends on connecting information across distinct experiences to derive knowledge that was never directly observed. Behavioral work indicates this capacity improves into emerging adulthood. However, it remains unclear whether these developmental gains reflect the quantitative strengthening of a single inference mechanism or qualitative changes in the knowledge representations and computations that support inference decisions. Here, we tested whether improvements in inference are linked to maturation of the hippocampus and posterior parietal cortex, resulting in age-related differences in how inference decisions are computed. We predicted that neural engagement in children (7–12 years) would track a variable-speed process, consistent with an iterative retrieval mechanism wherein distinct memories are retrieved and combined during inference. In contrast, neural engagement in adults would track a fixed-speed process, consistent with a direct inference mechanism supported by structured representations of shared relations across experiences. Using functional MRI combined with computational modeling of response times, we show that hippocampal activity predicts inference through iterative retrieval in both children and adults. In contrast, angular gyrus activity shows significant developmental differences in its relationship to both inference computations, with stronger associations with variable-speed processing in children and fixed-speed processing in adults. Moreover, we showed stronger relationships between fixed-speed computations and activation in both hippocampus and angular gyrus in adults relative to children. Together, these findings identify a developmental shift in the neural mechanisms supporting inference, suggesting that maturation of posterior parietal cortex may facilitate access to structured representations that capture links across events to enable more efficient inference.

## 1. Introduction

Inference allows us to discern information about the world that we have not directly experienced (Schlichting & Preston, 2015). This powerful cognitive ability is critical for new learning and adaptive decision making, as demonstrated by its association with academic achievement in children and college students (Esposito & Bauer, 2017; Varga et al., 2019). Understanding the mechanisms that support inference is therefore important for promoting educational outcomes across development. Although school age children are capable of inference (Bauer & Souci, 2010), research indicates a protracted developmental trajectory, such that even adolescents perform worse at inference than adults (Bauer, 2021; Schlichting et al., 2017; Shing et al., 2019). Importantly, while inference ability has been well characterized behaviorally across development, the neural computations that underlie developmental gains in reasoning are poorly understood. Here, we use computational modeling to quantify how the structure of memory representations impacts inference decisions across development. This approach moves beyond characterizing behavioral differences to identify the computations that support increasingly efficient inference with age.

One prominent suggestion is that children use different types of neural codes and computations to perform inference than adults (Bauer et al., 2021; Schlichting et al., 2022; Shing et al., 2019; Varga et al., 2025; Varga & Bauer, 2013). Specifically, adults may use knowledge they have already derived about event relationships to make inferences quickly and accurately. In contrast, children may rely on a less efficient mechanism, wherein they must retrieve and combine separate individual memories on the fly during inference. This difference in computation may make children’s inference decisions slower and potentially less accurate than those of adults. Neuroimaging work has implicated both the hippocampus and posterior parietal cortex in reasoning, particularly in adults (Crone et al., 2009; Morton et al., 2020; Wendelken, 2015; Zeithamova et al., 2012). However, most prior studies have focused on whether these regions are more or less engaged as a function of age or inference success, rather than on the specific computations they support during inference decisions. Here, we test the hypothesis that maturation of hippocampal and parietal systems gives rise to a qualitative shift in how inference is computed across development. We do so by leveraging a computational modeling approach that links trial-level neural responses to distinct inference mechanisms—one based on directly retrieving a derived relationship and another based on iteratively retrieving and combining individual memories online.

Let’s consider how one might achieve inference via two distinct mechanisms upon experiencing overlapping events that share a common feature. Imagine that an individual sees an acquaintance walking a dog at the park one day. A few days later, they see an unfamiliar man with the same dog at an outdoor cafe. Prior work has shown that adults can connect these two premise events in memory (Schlichting & Preston, 2015, 2016; Zeithamova et al., 2012; Zeithamova & Preston, 2010). Specifically, adults may integrate them through overlapping neural codes that promote the ability to directly infer connections between them (e.g., a relationship between the acquaintance and man based on their shared association with the dog; **Fig. 1A, middle panel**). Because adults have already integrated these events in memory, they may be able to directly retrieve and thus infer the “acquaintance-man” connection (i.e., *direct inference mechanism*; **Fig. 1B, middle panel**). Though previous work has not examined children’s neural codes during inference, behavioral studies indicate children are less likely and slower to make inferences (Bauer et al., 2021; Schlichting et al., 2017; Shing et al., 2019). Furthermore, children fail to use implicit connections between overlapping events to support their explicit inference judgments, in contrast to adults (Abolghasem et al., 2023).

**Figure 1.**
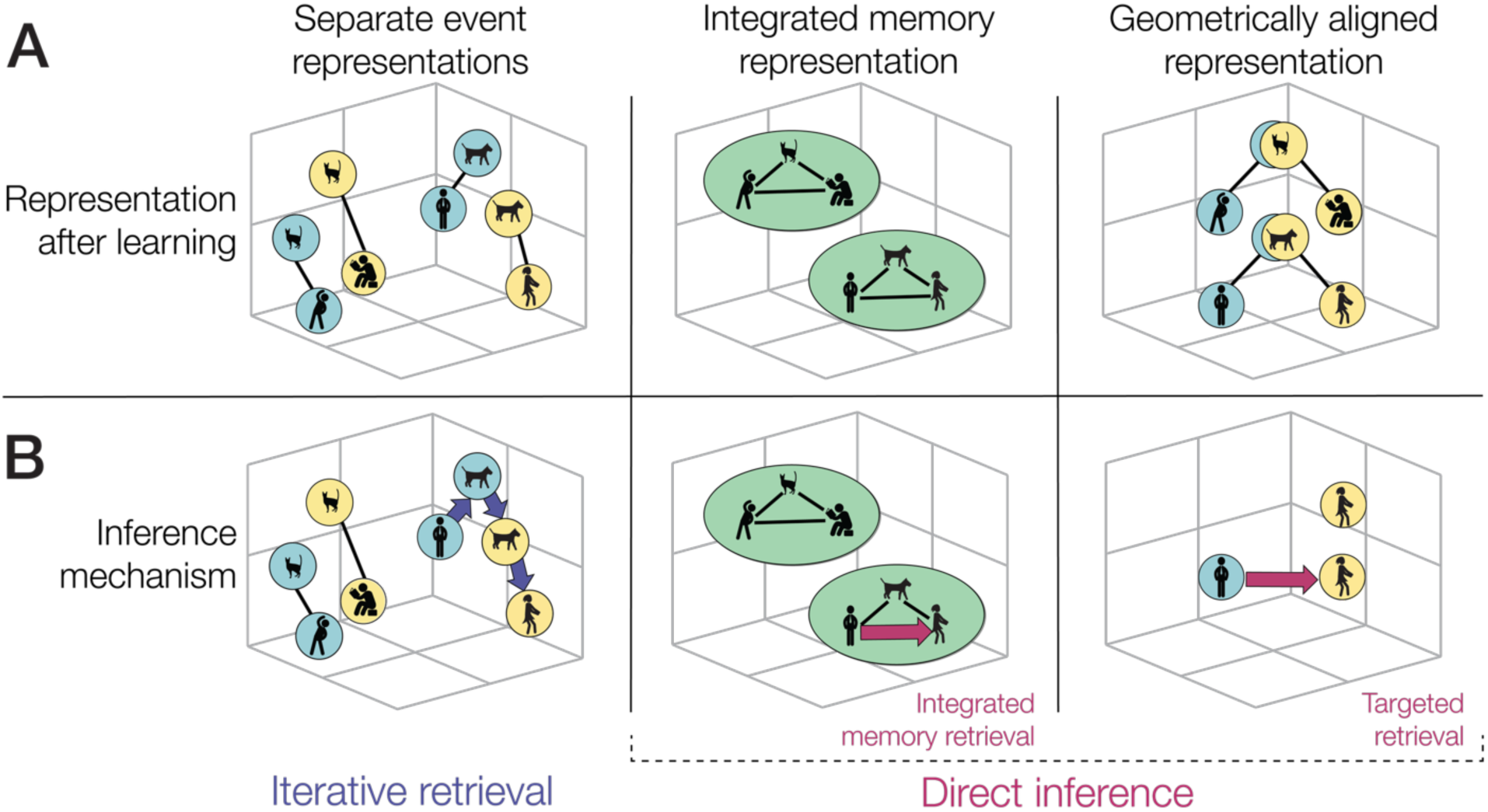
**(A)** After learning, overlapping experiences (e.g., AB and BC) can be represented in different ways within memory. **(B)** These different representations determine the computations supporting inference. When memories are stored as separate representations, inference requires iterative retrieval across associations (purple arrows; AB → BC; left panel). In contrast, when representations are integrated or geometrically aligned, the inferred relationship between premise pairs can be accessed more directly, supporting a direct inference computation (pink arrows; AC; middle and right panels). Because our approach cannot distinguish between integrated and geometrically aligned memory representations, we include both as potential accounts.

It’s been posited these developmental differences arise because children do not connect events as they are learned, but rather differentiate them as separate memory representations (Schlichting et al., 2022; Shing et al., 2019; Varga et al., 2025; **Fig. 1A, left panel**). This differentiation would require children to iteratively retrieve and combine individual events at the time of inference (Bauer et al., 2015, 2021; Shing et al., 2019; Varga & Bauer, 2013). For example, a child would need to iteratively retrieve the “acquaintance-dog” and “dog-man” premise events, and then combine them online to infer the acquaintance-man connection (i.e., *iterative retrieval mechanism*; **Fig. 1B**, **left panel**). This mechanism would explain the longer inference response times observed in children versus adults because it requires retrieving twice as many associations as the direct inference mechanism. The greater retrieval demands of iterative inference may also reduce accuracy, although both mechanisms can support successful inference. Prior behavioral and neuroimaging findings are consistent with a developmental shift in the mechanisms supporting inference (Bauer et al., 2015; Schlichting et al., 2022; Varga et al., 2025). However, prior studies have not directly examined the neural computations engaged at the time of inference decisions. Here, we combine computational modeling with fMRI to distinguish the mechanisms supporting successful inference in children and adults and how they shift during development.

Our computational model distinguishes inference mechanisms based on the number of associations that must be retrieved to reach a decision. The iterative retrieval mechanism requires retrieving multiple associations or premise pairs, whereas the direct inference mechanism requires retrieving only one association—the derived connection (Molitor et al., 2021; Morton et al., 2020). This difference between mechanisms should be reflected in how long it takes individuals to reach an inference decision. Inference should be slow if it requires iteratively retrieving multiple associations, and fast if it requires directly retrieving a single association.

We can therefore interrogate the predicted response time signatures of the proposed mechanisms by comparing inference decision speed with the time it takes for retrieval of individual premise pairs. Slower response time for inference versus premise pair retrieval would indicate a *variable speed process*, consistent with an iterative retrieval mechanism. This pattern would suggest that two individual associations must be retrieved and combined for inference. In contrast, similar response times for premise pair retrieval and inference would indicate a *fixed speed process*, consistent with a direct inference mechanism. This pattern would suggest that only one association is retrieved for inference. We can therefore model individual response times—across inference and premise pair tests—to identify which mechanisms contribute to inference. Application of this model to adult data suggests differential reliance on iterative retrieval and direct inference mechanisms contribute to variability in inference performance (Morton et al., 2020). Furthermore, distinct brain regions mediate these mechanisms in adults, with iterative retrieval supported by the hippocampus and direct inference by the lateral parietal cortex. We therefore use a similar approach to test whether the neural computations supporting inference differ across development and identify the brain regions associated with these differences.

Prior work has tied changes in brain function to the development of both memory (Ghetti & Bunge, 2012; Keresztes et al., 2017; Ofen et al., 2007) and reasoning behaviors (Crone et al., 2009; Modroño et al., 2018; Tooley et al., 2022; Wendelken et al., 2011). However, how the brain supports developmental gains in inference remains unclear, particularly at the time of computing inference decisions. One region we expect to be important to inference development is the hippocampus. Research indicates this region supports both differentiated and integrated memory representations in adults, depending on contextual demands (Chanales et al., 2017; Favila et al., 2018; Hulbert & Norman, 2015; Molitor et al., 2021; Schlichting et al., 2015a). Adults may therefore engage the hippocampus to support iterative retrieval when they possess differentiated representations and direct inference when they possess integrated representations. In contrast to adults, a recent study found that the hippocampus supported differentiated— but not integrated—memory representations in children (Varga et al., 2025). This finding aligns with work showing that children fail to reactivate related memories during encoding, precluding an opportunity for integration (Schlichting et al., 2022).

Together, these findings suggest the hippocampus may support only an iterative retrieval mechanism in children, wherein distinct memories are retrieved and combined at the time of inference. We therefore predict that the hippocampus supports inference across development, but does so through computations that vary by age. If true, hippocampal activation should predict response time patterns that are consistent with both iterative retrieval and direct inference mechanisms in adults, but only an iterative retrieval mechanism in children. Importantly, assessing hippocampal engagement during inference can only attest to the potential involvement of this region. Our unique combination of modeling and brain function is needed to tie this engagement to the hypothesized differences in underlying mechanism.

Beyond the hippocampus, the posterior parietal cortex has also been implicated in children’s emerging reasoning ability (Crone et al., 2009; Modroño et al., 2018; Tooley et al., 2022; Wendelken et al., 2011). However, we know little about the mechanisms whereby posterior parietal cortex may contribute to developmental improvements in inference. One possibility is that a mature posterior parietal cortex supports more accurate reinstatement of memory representations during inference decisions (Favila et al., 2018; Kuhl & Chun, 2014; Sestieri et al., 2017; Vilberg & Rugg, 2009). This possibility suggests posterior parietal cortex supports any inference mechanism operating on memory representations, regardless of whether those representations are integrated or differentiated. However, recent adult data suggest posterior parietal cortex may also support an emerging ability to navigate inferred shortcuts through learned memory representations (Morton et al., 2020). Through applying a computational model to adult neuroimaging data, researchers found that direct inference was associated with geometrically aligned neural representations in the posterior parietal cortex (Morton et al., 2020). These representations align shared relational structure across events, potentially providing more direct routes to inference. For example, the sequence of getting a basket, filling it with items, and waiting in the checkout line can be aligned across different grocery store trips (**Fig. 1A-B, right panel**). This work indicates that representations in the posterior parietal cortex likely support direct inference in adults.

Additional research indicates a specific posterior parietal cortex region, the angular gyrus, is potentially critical for supporting a shift to direct inference with age. The angular gyrus has been implicated in a variety of domain-general inference behaviors including arithmetic operations (Ischebeck et al., 2009), integration of spatial information (Sack, 2009), reading comprehension (Price & Mechelli, 2005), and attribution of mental states (Buckner et al., 2008). Moreover, there are protracted developmental changes in angular gyrus function and structure (Gogtay et al., 2004; Seghier, 2013; Westlye et al., 2010) that parallel developmental improvements in inference (Bauer, 2021; Schlichting et al., 2017; Shing et al., 2019). We therefore hypothesized that, within the posterior parietal cortex, angular gyrus would track a direct inference mechanism increasingly during development. Specifically, we predicted angular gyrus activation would predict response time patterns consistent with direct inference in adults but not children.

More broadly, these predictions reflect a central question about the computations that support inference. We will illustrate this question using letters that map onto our initial example. Specifically, you can think of the “acquaintance” as an A item, the “dog” as a B item, and the “unfamiliar man” as a C item. Now let’s say participants learned a series of premise AB and BC associations that overlap via a shared item, B. Is inference achieved by retrieving overlapping associations step-by-step (AB → BC), or by accessing direct links between indirectly related items from premise pairs (AC)? These alternatives make distinct predictions about both response time and neural engagement, providing a framework for testing how inference is computed across development. We assessed these predictions using model-based neuroimaging analyses. Children (7–12 years) and adult participants were asked to study non-overlapping and overlapping pairs of pictures (i.e., premise events). Overlapping pairs (AB, BC) consisted of one trial-unique picture and one picture shared with another pair, paralleling the structure of our “acquaintance–dog” and “dog–man” example. Participants were then scanned while being tested on their memory for overlapping and non-overlapping pairs (BC, XY; memory tests), as well as their ability to infer a connection between overlapping pairs (AC; inference test). Neuroimaging data were used to assess age-related differences in activation and to determine whether trial-by-trial variability in activation predicted response time patterns associated with iterative retrieval versus direct inference mechanisms. This approach allowed us to dissociate inference computations and identify how their underlying neural substrates change across development.

## 2. Method

### 2.1. Experiment overview

This experiment was part of a multi-session study conducted across three days (*M* delay: 4.8 weeks between days 1 and 2; 1.0 week between days 2 and 3). Here, we focus on data collected on day 3, when participants completed the associative inference task (see **Supplement** for details of days 1 and 2). The associative inference task is adapted from prior work in adults and children (Preston et al., 2004; Schlichting et al., 2022) and provides a controlled paradigm for assessing inference. Rather than relying on real-world events, the task uses arbitrary but familiar visual stimuli to isolate the memory processes that support inference.

### 2.2. Participants

Seventy-seven participants completed the multi-session study through day 3. Three participants were excluded due to a technical problem with the scanner (n = 2) or noncompliance (n = 1), resulting in a final sample of 74 participants. This sample included 50 children aged 7–12 years (*M* = 10.10, SEM = 0.23; 6–9 children per year; 27 female) and 24 young adults aged 19–33 years (*M* = 26.99, SEM = 0.85; 12 female). Additional partial data exclusions were made for neuroimaging analyses and are described in detail in the relevant sections below. To preview, ten additional participants were excluded from the univariate GLM analysis due to a lack of usable scan data for at least one experimental condition, reducing the sample to 64 participants (41 children and 23 young adults). A follow-up analysis of angular gyrus engagement excluded an additional participant due to their extreme parameter estimates, reducing the sample to 63 participants (41 children and 22 young adults). No participants were excluded from the computational response-time analysis given its ability to accommodate missing data.

Participants had average or above-average IQ (scored no lower than 2 SD below the mean on the Wechsler Abbreviated Scale of Intelligence, Second Edition; Wechsler, 2011) and psychological symptoms within the typical range (if child: scored below the clinical range on the Child Behavior Checklist; Achenbach, 1991; if adult: scored below 1 SD above the normative sample mean on the Symptom Checklist 90-Revised; Derogatis, 1977). All participants had no reported developmental or psychological disorders and were native English speakers (learned by age 3).

The sample was 6.8% Asian, 6.8% Black, 77.0% White, and 9.5% multiracial; 12.2% also identified as Hispanic. Annual household income was assessed for all members of the household (if child) or for parents only (if adult) and reported for 66 participants (89.2% of the sample). Of these participants, 6.1% had an annual household income below $40,000 USD, 51.5% between 40,000-150,000 USD, and 42.4% above 150,000 USD. Highest parental education level was reported for 73 participants (98.6% of the sample). Of these participants, highest parental education level was a high school diploma or its equivalent for 2.7%, vocational/technical school or college without receiving a bachelor’s degree for 13.7%, a bachelor’s degree for 21.9%, and a graduate degree for 61.6%. The majority of participants were therefore of middle- to high-socioeconomic status. This distribution is representative of families who volunteer for developmental research through community outreach in Austin, Texas.

The study protocol was approved by the Institutional Review Board of the University of Texas at Austin. Informed consent was obtained from adult participants, and parental consent and child assent were obtained for child participants. Participants received monetary compensation ($25/hour) and a small prize (e.g., Pokémon cards, lip gloss, or gift cards valued at $5 or less).

#### 2.2.1. Design Considerations and Sample Size

An a priori power analysis was not conducted. Instead, we aimed for a sample size consistent with prior studies on the development of memory-based inference that used similar paradigms, which included 20-25 participants per age group (Schlichting et al., 2022; Shing et al., 2019). Our final sample size is also consistent with the only functional neuroimaging study of memory-based inference development, which included a minimum of 21 participants per age bin based on a power analysis indicating this number would result in 80% power to detect a behavioral effect of a within-subject manipulation of integration between groups (Schlichting et al., 2022). For recruitment purposes only, we divided our sample into three age bins of 7–9 years, 10–12 years, and young adults, aiming for approximately 20-25 participants within each bin, consistent with this prior work. However, all analyses examined age continuously, resulting in a total number of child participants that exceeds this reference point.

### 2.3. Materials

#### 2.3.1. Stimuli selection

We equated stimulus familiarity across participants, given evidence that less familiar stimuli are more difficult to integrate for knowledge generation (Reder et al., 2015). Stimuli included 40 images of faces and scenes (20 each) from popular childhood movies and television shows, as well as 120 common objects. To control for variability in exposure to the content from movie and television shows, participants completed a “know game” on day 1 to generate a personalized set of familiar faces and scenes (taken from Schlichting et al., 2022). During this task, participants viewed up to 225 images of characters (e.g., Aladdin) and scenes (e.g., Cinderella’s castle) presented in random order. For each image, they rated familiarity (“not at all familiar,” “a little bit familiar,” or “very familiar”) and provided a name or description if they reported any familiarity. Experimenters scored descriptions online as accurate or inaccurate to verify familiarity. We used these ratings to select 40 highly familiar and accurately identified faces and scenes for each participant. If fewer than 40 images met both criteria, we included items rated as less familiar but accurately identified. To ensure stimulus diversity, we included only one image from each movie or television show. The task terminated once participants identified 40 qualifying stimuli, and most participants did not view all 225 images.

We used a common set of 120 objects across participants, as exposure to these items is expected to be less variable. We selected objects that were easily nameable and broadly familiar across age groups based on age-of-acquisition norms (Kuperman et al., 2012). We randomly assigned objects to conditions for each participant.

#### 2.3.2. Organization of stimuli into picture triads and pairs

We combined 40 face and scene images with 80 object images to form 40 triads, each consisting of a face or scene (A item) paired with two objects (B and C items). This yielded 20 face-object-object (A_F_B_O_C_O_) triads and 20 scene-object-object (A_S_B_O_C_O_) triads. From these triads, we generated 40 initial pairs (AB) and 40 overlapping pairs (BC), such that each pair type shared a common B item. We combined the remaining 40 objects into 20 non-overlapping object–object pairs (XY). These pairs allowed us to compare performance for associations that shared content (overlapping pairs) versus those that did not (non-overlapping pairs). We included fewer non-overlapping pairs based on pilot testing, which showed that longer task variants (with equal numbers of pair types) increased fatigue and reduced performance, particularly in younger children. This design kept task duration manageable while maximizing power to assess inference across overlapping pairs.

### 2.4. Associative inference paradigm

#### 2.4.1. Phase 1: Initial pair learning

Participants first completed the initial pair learning phase, which consisted of study-test cycles (**Fig. 2A**). During each study phase, participants viewed 40 initial pairs (AB) sequentially for 3.5 s, separated by a 0.5 s fixation cross (B item on the right). Participants were instructed to generate a story linking the two images in each pair and to use any additional strategies to support memory. These instructions encouraged attention to both items and promoted consistent encoding strategies across ages.

**Figure 2.**
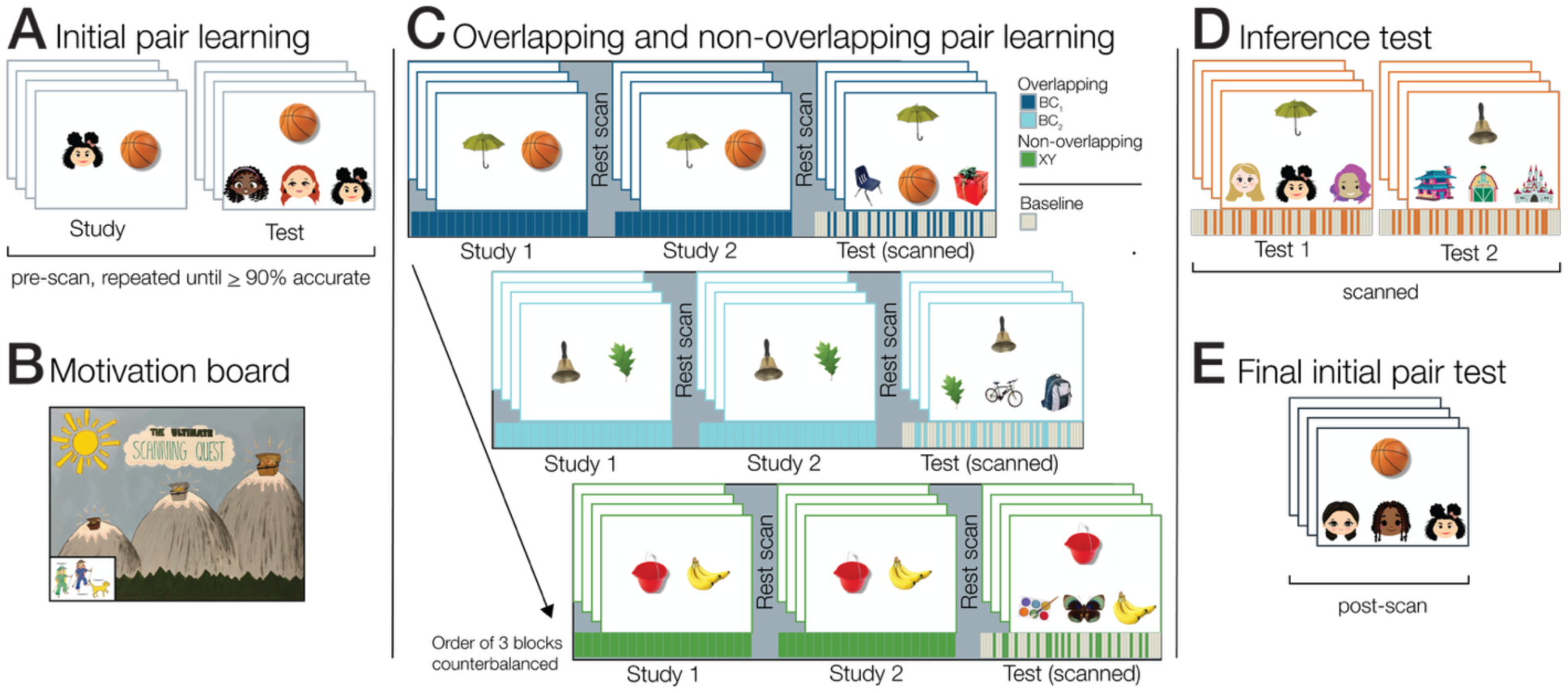
Associative inference task design. The faces and scenes in the real task were from popular childhood movies and television shows; they are replaced with images generated with DALL-E here due to copyright protections. **(A)** Participants first learned initial associations (AB) through study–test cycles prior to scanning. **(B)** A motivation board tracked progress toward a performance-based bonus during the scanning session. **(C)** In the scanner, participants learned overlapping pairs (BC_1_, BC_2_) and non-overlapping pairs (XY) across three blocks. Each block included study phases followed by a scanned test phase. Overlapping pairs shared elements with the initial AB pairs, providing the basis for later inference, whereas non-overlapping pairs served as a control. Test trials were jittered with baseline trials. **(D)** Participants then completed a surprise inference test (AC), requiring them to infer indirect relationships between items that had never been presented together (i.e., linking A and C through prior AB and BC learning). On each trial, participants selected the “distant friend” (A) associated with a cue (C). **(E)** Finally, participants completed a post-scan test of initial pairs (AB) to assess memory for the initial associations.

Participants then completed a self-paced test phase assessing memory for the initial (AB) pairs using a three-alternative forced choice format. On each trial, participants viewed a B item cue and selected the associated A item from three options presented below. Response options included the target A item and two lure A items drawn from other pairs and matched in category (face or scene). Participants responded using a button box (1 = left, 2 = middle, 3 = right). After each response, the lure items disappeared, leaving the cue and target on screen, followed by 1 s of feedback. Correct responses triggered the message “*Right! The answer is:*” in green text with a positive tone, whereas incorrect responses triggered “*Wrong. The answer is:*” in red text with a negative tone.

At the end of each test run, participants received summary feedback based on accuracy. Participants whose accuracy was ≤ 34% for that run saw “*That’s okay. Let’s try harder!*” and the outline of three empty stars. Those with higher accuracy saw more positive text and rainbow stars. Accuracy between 35-59% resulted in “*Good job!*” and one rainbow star; accuracy between 60-84% resulted in “*Great job!*” and two rainbow stars; and accuracy ≥ 85 resulted in “*Awesome job!*” and three rainbow stars. Together, trial- and test-level feedback encouraged pair learning and task engagement. Participants completed study-test cycles (with trial order randomized each cycle) until they had completed at least two cycles and demonstrated high memory for the initial pairs (i.e., 90-100% accuracy). This procedure ensured all participants formed robust initial pair memories.

#### 2.4.2. Phase 2: Overlapping and non-overlapping pair learning

Participants next completed overlapping and non-overlapping pair learning in three blocks (**Fig. 2C**). In each block, participants learned 20 pairs. Two blocks contained overlapping pairs (BC_1_, BC_2_), and one block contained non-overlapping pairs (XY). Block order was fully counterbalanced across participants. Critically, we did not inform participants about the overlap between initial and overlapping pairs, nor did we indicate that they would later need to make inferences across them.

To support engagement during this longest phase of the task, participants completed this phase in conjunction with a motivation board (**Fig. 2B**). Before entering the scanner, the experimenter introduced a board depicting three mountains with a treasure chest at each peak, increasing in size from left to right. Each mountain corresponded to one block of overlapping or non-overlapping learning. Participants advanced a character piece across the mountains after each block and received a brief break (approximately 5 minutes) outside the scanner to move their piece after blocks one and two. At the end of the study, participants selected a prize (value < $5) from a prize box corresponding to the highest mountain they reached. This procedure encouraged engagement, provided regular scanning breaks, and allowed participants to track their progress.

Each block consisted of two study phases followed by a test phase (study phase 1, study phase 2, test phase). The two study phases were identical with the exception that trial order was randomized within each. The study phases were not scanned and repeated to promote robust learning of overlapping and non-overlapping pairs. A 5 min 10 s resting-state scan followed each study phase. These scans are not analyzed here and reported only to depict the full experimental procedure.

Study phase trials followed the same structure as in initial pair learning, with different stimuli. Participants again viewed pairs sequentially for 3.5 s with a 0.5 s fixation cross ISI (B and Y items on the right). For each pair, participants were instructed to generate a story linking the two images and to use any additional strategies to support memory.

We scanned the test phases. Each test phase consisted of three-alternative forced choice trials in which participants viewed a cue item (C for overlapping pairs; X for non-overlapping pairs) and selected the associated item. For overlapping pairs, response options included the target B item and two lure B items drawn from other overlapping pairs. For non-overlapping pairs, response options included the target Y item and two lure Y items drawn from other non-overlapping pairs. Participants had up to 8 s to respond using a button box (1 = left, 2 = middle, 3 = right). If participants did not respond within 6 s, a red border appeared to prompt a response within the remaining 2 s. After a response, a gray border appeared to indicate that the response was registered.

Trials were separated by a 4 s ISI, and we interspersed 30 baseline trials to provide jitter. Baseline trials lasted 2 s (including a 0.5 s ISI) and required participants to indicate whether a dot appeared in a left, center, or right box using the same response mapping. We added four additional baseline trials to the beginning and end of each test phase to account for stabilization and lag of the hemodynamic response function during scanning. Participants received trial-level and run-level feedback using the same procedures and performance thresholds described in initial pair learning.

#### 2.4.3. Phase 3: Inference test

Participants were then scanned while completing a surprise inference test that assessed their ability to infer relationships between initial (AB) and overlapping pairs (BC; **Fig. 2D**). The test included 40 inference (AC) trials (one per triad), which we divided into two runs of 20 trials to maintain reasonable scan durations. The same trial timing and structure as in overlapping and non-overlapping pair tests were used, including baseline trials and performance feedback at the end of each trial and run. However, the task differed in the stimuli and the relation being tested. Each inference trial followed a three-alternative forced choice format in which participants viewed a C item and selected which of three response options it shared an indirect relationship with. Response options included the target A item from the same ABC triad (linked via the shared B item) and two lure A items drawn from other triads.

Before the inference test, the experimenter explained the indirect AC relationship using a “distant friendship” analogy to facilitate understanding. While displaying an example initial and overlapping pair, the experimenter highlighted that the A and C items shared a common associate (the B item), making them “distant friends.” Participants were instructed to identify these distant friends during the test. The experimenter did not begin the test until participants demonstrated understanding by correctly answering a practice trial and explaining the task in their own words.

Most participants completed both inference test runs in the scanner. However, due to scan time constraints, a subset completed one run (n = 8; three adults and five children) or both runs (n = 6; one adult and five children) outside the scanner. Those who completed both runs outside the scanner were necessarily excluded from the GLM analysis due to a lack of scan data for the inference condition, but were included in the computational response-time analysis given its ability to accommodate missing data.

#### 2.4.4. Phase 4: Final test of initial pair memory

Participants completed a final test of initial pairs (AB) during the last task phase outside of the scanner (**Fig. 2E**). We administered this test to ensure that poor inference performance could not be attributed to insufficient memory for the initial pairs. The test was identical to that used during initial pair learning (with trial order randomized) but was administered only once.

### 2.5. MRI data acquisition

We acquired MRI data on a 3 Tesla Siemens Skyra scanner in the Biomedical Imaging Center (RRID: SCR_021728), a core facility within the Center for Biomedical Research Support at the University of Texas at Austin. High-resolution whole-brain functional data were collected using a T2*-weighted multiband-accelerated EPI sequence (TR = 2000 ms, TE = 30 ms, flip angle = 77°, FOV = 220 mm, matrix = 128 × 128 × 78, 1.7 mm isotropic voxels, multiband factor = 3, GRAPPA factor = 2, phase partial Fourier = 7/8). A field map (TR = 668 ms, TE = 5 ms / 7.46 ms, flip angle = 5°, FOV = 192 mm, matrix = 128 × 128 × 67, 1.5 × 1.5 × 2 mm voxels) was also acquired each time participants re-entered the scanner (e.g., following a break) or when head position appeared to shift, to correct for distortion and blurring due to magnetic field inhomogeneities.

We acquired a high-resolution whole-brain T1-weighted 3D MPRAGE (TR = 1900 ms, TE = 2.43 ms, TI = 900 ms, flip angle = 9°, FOV = 256 mm, 192 slices, 1 mm isotropic voxels, GRAPPA factor = 2) on day 3 for all participants, as well as on day 2 for all but three participants. Most participants thus had two T1w volumes. When time permitted, we also acquired a T2-weighted FLAIR image (TR = 5000 ms, TE = 397 ms, TI = 1800 ms, FOV = 256 mm, 192 slices, 1 mm isotropic voxels, slice partial Fourier = 7/8, GRAPPA factor = 2) on day 2. One to two coronal T2-weighted structural images oriented perpendicular to the hippocampal axis (TR = 13150 ms, TE = 82 ms, matrix = 384 × 60 × 384, 0.4 × 0.4 mm in-plane resolution, 1.5 mm slice thickness, 60 slices, flip angle = 150°, FOV = 166 mm) were additionally collected on day 2. Our focus is on the day 3 data. However, when obtained, the high-resolution whole-brain T1-weighted 3D MPRAGE from day 2 and the T2-weighted FLAIR image from day 2 were also used to improve co-registration and normalization of functional data to MNI space and cortical surface reconstruction. The coronal T2-weighted structural images are not analyzed here.

### 2.6. MRI data processing

We preprocessed MRI data using a pipeline developed by Morton (2022) that incorporates FMRIB Software Library version 5.0 (FSL; Smith et al., 2004) and Advanced Normalization Tools (ANTs; Avants et al., 2011). Each high-resolution T1-weighted (T1w) 3D MPRAGE first underwent bias-field correction using N4BiasFieldCorrection (Tustison et al., 2010). For participants with two anatomical scans, we created a common anatomical space by co-registering the two T1w volumes using the nonlinear antsRegistration tool (Avants et al., 2008), scaling them using a single multiplicative factor to match intensity, and averaging the resulting volumes.

We then performed cortical reconstruction using recon-all in FreeSurfer (Dale et al., 1999) on either the averaged T1w volume (for participants with two scans) or the single T1w volume (for participants with one scan). When available, the T2-weighted FLAIR image was used to refine cortical surface reconstruction via enhanced pial extraction. FreeSurfer outputs were registered back to the original T1w space using antsRegistration (Avants et al., 2008) to enable their use in functional data processing.

We preprocessed functional time series data by applying bias-field correction (N4BiasFieldCorrection; Tustison et al., 2010), motion correction using spline interpolation in MCFLIRT (Jenkinson et al., 2002), and skull stripping using BET (Smith, 2002). Quality control and motion scrubbing were conducted using the fmriqa package (Poldrack et al., 2013). We further applied field map-based distortion correction by first bias-field correcting the field map magnitude image (N4BiasFieldCorrection; Tustison et al., 2010) and then registering it to the temporally closest T1w volume using antsRegistration (Avants et al., 2008). FSL’s epi_reg tool was then used to simultaneously unwarp the functional data and perform boundary-based registration to this T1w volume. This registration was refined over 20 iterations using antsRegistration (Avants et al., 2008), which updated an unwarped functional target image at each iteration using FUGUE (Jenkinson, 2004). Next, we nonlinearly registered the average unwarped functional images to a reference functional run (the second test run during the overlapping and non-overlapping learning phase) using antsRegistration (Avants et al., 2008). After estimating all transformations, we concatenated motion correction, unwarping, and registration steps and applied them to the raw functional data using B-spline interpolation in two steps to minimize interpolation error. Runs were excluded from subsequent analyses if they showed excessive motion, defined as more than one-third of volumes with mean framewise displacement > 0.30 mm.

### 2.7. Regions of interest (ROIs)

We focused neuroimaging analyses on whole-brain gray matter, the hippocampus, and the angular gyrus (with additional exploratory regions described in the **Supplement**). We defined ROIs in each participant’s native T1w space using FreeSurfer atlases. These native-space masks were used for computational response-time model analyses. We also generated group-level ROI masks for univariate GLM analyses. To do so, we warped each participant’s native-space masks to 1 mm standard MNI space using nearest-neighbor interpolation with antsRegistration (Avants et al., 2008). We then averaged the warped masks to create group probability maps for each ROI. Finally, we thresholded these maps at 0.50 to include voxels labeled as part of the ROI in at least 50% of participants.

### 2.8. Behavioral analyses

Behavioral analyses interrogated age-related differences in memory and inference performance. All analyses were conducted in R (R Core Team, 2026). The lme4 package (Bates et al., 2015) was used to run models. The stats, easystats (Ludecke et al., 2022), car (Fox & Weisberg, 2019), matrix (Bates et al., 2025), plotrix (Lemon, 2006), modelbased (Makowski et al., 2025), and reports (Makowski et al., 2025) packages were used for additional statistics and reporting assistance, and the ggplot2 package (Wickham, 2016) for data visualization. Data were analyzed using a model comparison approach with nested linear or linear mixed-effects models. We fit two models for each analysis: a main effects model that included all predictors of interest, and an interaction model that included the same predictors along with their interactions with age to test developmental hypotheses. Age was always calculated in years and mean-centered (see **Supplement** for models that instead assessed age group differences between children and adults). Categorical variable effects were always examined using deviation coding, such that comparisons were made to the grand mean instead of a specified reference level. Model fit was compared using a likelihood ratio chi-square test. Results from the better-fitting model are reported. For the linear mixed-effects models, results include fixed-effect estimates and their significance based on Wald chi-square tests.

#### 2.8.1. Analysis of initial pair memory

We used nested linear mixed-effects models to assess whether participants: (1) formed robust memories for initial (AB) pairs (**Fig. 2A**) and (2) maintained these memories across the task, as indexed by performance on the final initial pair test (**Fig. 2E**). The main effects model included age and test phase (initial pair learning vs. final test) as fixed effects and participant as a random intercept (accuracy ∼ age + test phase + (1 | participant)). The interaction model included an additional age × test phase interaction to assess developmental differences in retention (accuracy ∼ age + test phase + age × test phase + (1 | participant)).

During initial pair learning, participants completed study-test cycles until they completed at least two cycles and reached high accuracy (90-100%). All but one adult participant met this criterion within two cycles (group-level mean accuracy: cycle 1 = 94%, cycle 2 = 98%). One adult required four cycles but performed above chance throughout (cycle 1 = 78%, cycle 2 = 85%, cycle 3 = 88%, cycle 4 = 100%). We therefore quantified initial pair learning as accuracy during cycle 2 for all participants except this individual, for whom we used cycle 4. We assessed final test performance using the single administration of the final initial pair test.

We expected high accuracy overall, with slightly lower performance on the final test compared to initial learning, reflecting modest memory decay. Although participants across ages achieved high accuracy during learning, we also tested whether children showed lower initial performance and/or greater decay over time. A likelihood ratio chi-square test indicated that the interaction model did not improve fit relative to the main effects model (χ^2^(1) = 1.81, *p* = 0.178). We therefore report results from the main effects model.

#### 2.8.2. Analysis of memory versus inference performance

We used two sets of nested linear mixed-effects models to compare performance across non-overlapping memory (XY), overlapping memory (BC), and inference (AC) tests (**Fig. 2C-D**). One set modeled response accuracy, and the other modeled response time for correct trials only. For each outcome, the main effects model included age and test type (non-overlapping memory, overlapping memory, inference) as fixed effects and participant as a random intercept (performance ∼ age + test type + (1 | participant)). The interaction model additionally included an age × test type interaction (performance ∼ age + test type + age × test type + (1 | participant)).

We predicted a main effect of test type, with better performance—higher accuracy and faster response times for correct trials—on the memory tests compared to the inference test. We also predicted an age × test type interaction. Although performance should improve with age overall, we expected more protracted improvement for inference, reflecting a potential shift from iterative retrieval during childhood to more efficient, direct inference in adulthood.

Likelihood ratio chi-square tests indicated that the interaction model provided a better fit than the main effects model of accuracy (χ^2^(2) = 19.58, *p* < 0.001), but not of response time (χ^2^(2) = 0.04, *p* = 0.978). We therefore report results from the interaction model for accuracy and the main effects model for response time. Given performance similarities between non-overlapping and overlapping memory tests, we collapsed these conditions into a single memory condition in all subsequent analyses.

### 2.9. Neuroimaging analyses

Neuroimaging analyses were organized around three successive analytical goals, which we overview here. First, we used a univariate GLM analysis to identify brain regions that were differentially engaged during successful inference relative to premise pair retrieval and determine whether their recruitment varied across development. This analysis allowed us to establish whether children and adults engage the same neural systems when retrieving learned associations versus inferring relationships that were never directly experienced, providing an initial characterization of developmental differences in the neural systems supporting inference.

Second, we sought to determine whether a region that was differentially engaged by children and adults for inference relative to premise pair memory tracked behavioral performance. Specifically, we tested whether this region was associated with faster successful inference decisions and whether this relationship differed between children and adults. This analysis allowed us to move beyond identifying developmental differences in neural recruitment to test if neural engagement was associated with the efficiency of inference.

Our third analysis expanded upon the second by using a computational response-time model to distinguish between two mechanisms that can support successful inference: an iterative retrieval mechanism, whereby individual memories are retrieved and combined at the time of inference, and a direct inference mechanism, in which previously derived relationships can be accessed more efficiently. These proposed mechanisms make distinct predictions about response times during premise pair retrieval and inference, allowing us to model their contributions as variable-speed and fixed-speed processes, respectively. By incorporating trial-level neural activation into the model, we could then determine how neural engagement during inference related to these distinct computations and whether those relationships changed across development. Thus, this final analysis allowed us to test whether developmental improvements in inference reflect more efficient operation of the same retrieval mechanism or differences in the neural computations supporting inference decisions. The details of these analytical strategies can be found in the immediately following sections.

#### 2.9.1. Univariate GLM analysis

##### 2.9.1.1. Neural activation during memory versus inference

We conducted a univariate GLM analysis to quantify differences in neural activation during memory and inference. Specifically, we tested whether regions showed greater engagement during successful inference (inferring a connection among two associations, i.e., AC) than during successful memory (retrieval of a single association, either BC or XY), and whether this difference varied with age. We predicted greater hippocampal engagement for inference than memory across all ages, consistent with a role in supporting both iterative retrieval and direct inference. In contrast, we predicted an age-related increase in posterior parietal cortex engagement for inference relative to memory, reflecting its proposed role in more efficient, direct inference (Morton et al., 2020) and its protracted developmental trajectory (Gogtay et al., 2004; Westlye et al., 2010).

##### 2.9.1.2. Partial data exclusions

We excluded runs from this analysis if they showed excessive motion, defined as more than one-third of volumes with mean framewise displacement > 0.30 mm, or if they were not acquired due to scanner time constraints or technical difficulties. These criteria resulted in ten participants who lacked usable scan data for at least one experimental condition. These participants were therefore excluded from the analysis, yielding a final sample of 64 participants: 18 young children (7–9 years; *M* = 8.79 years; SEM = 0.15; 10 female), 23 older children (10–12 years; *M* = 11.49 years; SEM = 0.20; 12 female), and 23 young adults (19–33 years; *M* = 26.78 years; SEM = 0.86; 11 female). Age bins are reported for descriptive purposes only; all analyses treated age as a continuous variable.

##### 2.9.1.3. Statistical modeling

We analyzed functional data using the fMRI Expert Analysis Tool (FEAT; Woolrich et al., 2001, 2004) in the FMRIB Software Library (FSL; Smith et al., 2004). A general linear model (GLM) was specified based on the experimental design and applied to the preprocessed functional data across three analysis levels.

First-level analyses modeled trial-level activity within each of the five test runs: the two overlapping memory test runs (BC), single non-overlapping memory test run (XY), and two inference test runs (AC). The model included four regressors: (1) correct trials (primary regressor of interest), (2) incorrect trials, (3) response time for correct trials, and (4) response time for incorrect trials. We modeled each trial using its onset, duration (8 s), and weight (1 for both correct and incorrect trials; mean-centered response time for response-time regressors). We included additional confound regressors to account for residual motion and added temporal derivatives to accommodate small timing shifts. The full model was convolved with a double-gamma hemodynamic response function, and spatial smoothing (FWHM = 4 mm), high-pass temporal filtering (cutoff = 128 s), and FILM prewhitening were applied. We then warped resulting contrast of parameter estimate images (COPES) to 1 mm MNI space and resampled them to functional resolution using antsRegistration (Avants et al., 2008), placing all data in a common space for higher-level analyses.

Second-level analyses combined runs within each participant using fixed-effects modeling. Inputs were the warped COPES for correct trials from each test run. The two overlapping memory test runs (BC) and single non-overlapping memory test run (XY) were grouped into a memory condition. The two inference test runs (AC) were grouped into an inference condition. We then computed our contrast of interest, inference > memory. This analysis yielded a mean COPE image for each participant.

Third-level analyses examined group-level effects using mixed-effects modeling implemented in FMRIB’s Local Analysis of Mixed Effects (FLAME 1). Inputs were participant-level COPE images from the second-level analysis. The model included two regressors of interest: mean activation (group-level effect of the contrast) and age (to assess developmental change). We included two additional confound regressors to control for response time differences: (1) mean response time across correct trials and (2) the difference between mean response time for inference and memory trials. We mean-centered age and response time regressors prior to analysis. This analysis produced a statistical image for each regressor of interest.

We identified significant clusters using Analysis of Functional NeuroImages (AFNI) functions (Cox, 1996; Cox et al., 2017a, 2017b). Spatial smoothness of residual noise within gray matter (whole-brain correction) and hippocampus (a priori ROI) was first estimated using 3dFWHMx with the spatial autocorrelation function. These estimates were used in 3dClustSim to generate 10,000 null datasets. Based on these simulations, we determined the minimum cluster size required to achieve a cluster-wise significance threshold of *p* < .05 following voxel-wise thresholding at *p* < .01 (two-sided, second-nearest neighbor clustering). The resulting thresholds were 956 voxels for gray matter and 96 voxels for hippocampus. Clusters exceeding these thresholds are reported. Mango was used to create a surface rendering of the brain when needed for visualization purposes, for this and all other analyses (Research Imaging Institute, UTHSCSA, n.d.)

#### 2.9.2. Interrogating angular gyrus engagement

Whole-brain analyses identified an age-related increase in posterior parietal cortex engagement for successful inference relative to memory. Within this region, the angular gyrus was an *a priori* region of interest based on the hypothesis that it supports a direct inference mechanism that emerges with development. To test this hypothesis, we examined angular gyrus engagement during successful inference relative to memory and its relationship to inference performance. Because accuracy approached ceiling in adults (*M* = 92.45%, SEM = 2.23%; **Fig. 4B**), we focused behavioral analyses on mean response time for correct inference trials. If the angular gyrus supports a more efficient, direct inference mechanism with age, then greater recruitment of this region during successful inference should be associated with faster correct inference, particularly in adults.

##### 2.9.2.1. Partial data exclusions

We excluded the same ten participants removed from the univariate GLM analysis due to a lack of usable scan data in at least one experimental condition. We additionally excluded an adult participant with extreme angular gyrus parameter estimates that aligned with experimental predictions (> 7 SD above the mean for inference and below the mean for memory), to ensure their data did not overly influence group-level effects. The final sample for this analysis therefore included 63 participants: 18 young children (7–9 years; *M* = 8.79 years; SEM = 0.15; 10 female), 23 older children (10–12 years; *M* = 11.49 years; SEM = 0.20; 12 female), and 22 young adults (19–33 years; *M* = 26.89 years; SEM = 0.89; 11 female). Age bins are reported for descriptive purposes only; all analyses treated age as a continuous variable.

##### 2.9.2.2. Extracting and analyzing angular gyrus parameter estimates

We extracted angular gyrus parameter estimates (PEs) for successful inference and memory for each participant. To control for response time differences between conditions, we re-estimated the univariate GLM with average response time for correct trials included as a nuisance regressor at the second level (rather than at the third level, as in the primary analysis). We then applied a structural angular gyrus mask to the second-level statistic images for memory and inference. We computed the mean signal within this mask to obtain angular gyrus PEs for correct memory and correct inference for each participant, which were standardized within participant. Similar to the behavioral analyses, these PEs were entered into two sets of nested models in R (R Core Team, 2026) using the lme4 package (Bates et al., 2015). The stats, easystats (Lüdecke et al., 2022), car (Fox & Weisberg, 2019), matrix (Bates et al., 2025), plotrix (Lemon, 2006), modelbased (Makowski et al., 2025), emmeans (Lenth & Piaskowski, 2026), marginaleffects (Arel-Bundock et al., 2024), checkmate (Lang, 2017), and reports (Makowski et al., 2023) packages were used for additional statistics and reporting assistance, and the ggplot2 package (Wickham, 2016) to visualize data.

We first tested whether age and test type (memory, inference) predicted angular gyrus engagement using linear mixed-effects models. The main effects model included age and test type as fixed effects and participant as a random intercept (angular gyrus PE ∼ age + test type + (1 | participant)). The interaction model additionally included an age × test type interaction (angular gyrus PE ∼ age + test type + age × test type + (1 | participant)). A likelihood ratio chi-square test indicated that the interaction model provided a better fit than the main effects model (χ^2^(1) = 6.09, *p* = 0.014). We therefore report results from the interaction model.

We next tested whether age and angular gyrus engagement during inference predicted response time for correct inference trials using linear models. The main effects model included age and angular gyrus PE as predictors (response time ∼ age + angular gyrus PE). The interaction model additionally included an age × angular gyrus PE interaction (response time ∼ age + angular gyrus PE + age × angular gyrus PE). An ANOVA indicated that the interaction model provided a better fit than the main effects model (*F*(1, 59) = 6.18, *p* = 0.016). We therefore report results from the interaction model.

#### 2.9.3. Computational response-time model analysis

Our univariate GLM analysis allowed us to examine associations between neural engagement and response time behavior, but not whether hippocampus and angular gyrus activation relate to distinct inference computations. To address this question, we fit response times using a Linear Ballistic Accumulator (LBA) model (Brown & Heathcote, 2008; Molitor et al., 2021; Morton et al., 2020) within a hierarchical Bayesian framework. The model formalized two candidate inference mechanisms. Iterative retrieval was implemented as a variable-speed process, producing slower responses for inference than memory because it requires retrieval of two associations (**Fig. 3A-B**). Direct inference was implemented as a fixed-speed process, producing similar response times for memory and inference because integrated or geometrically aligned representations support one-step inference (**Fig. 3A,C**).

**Figure 3.**
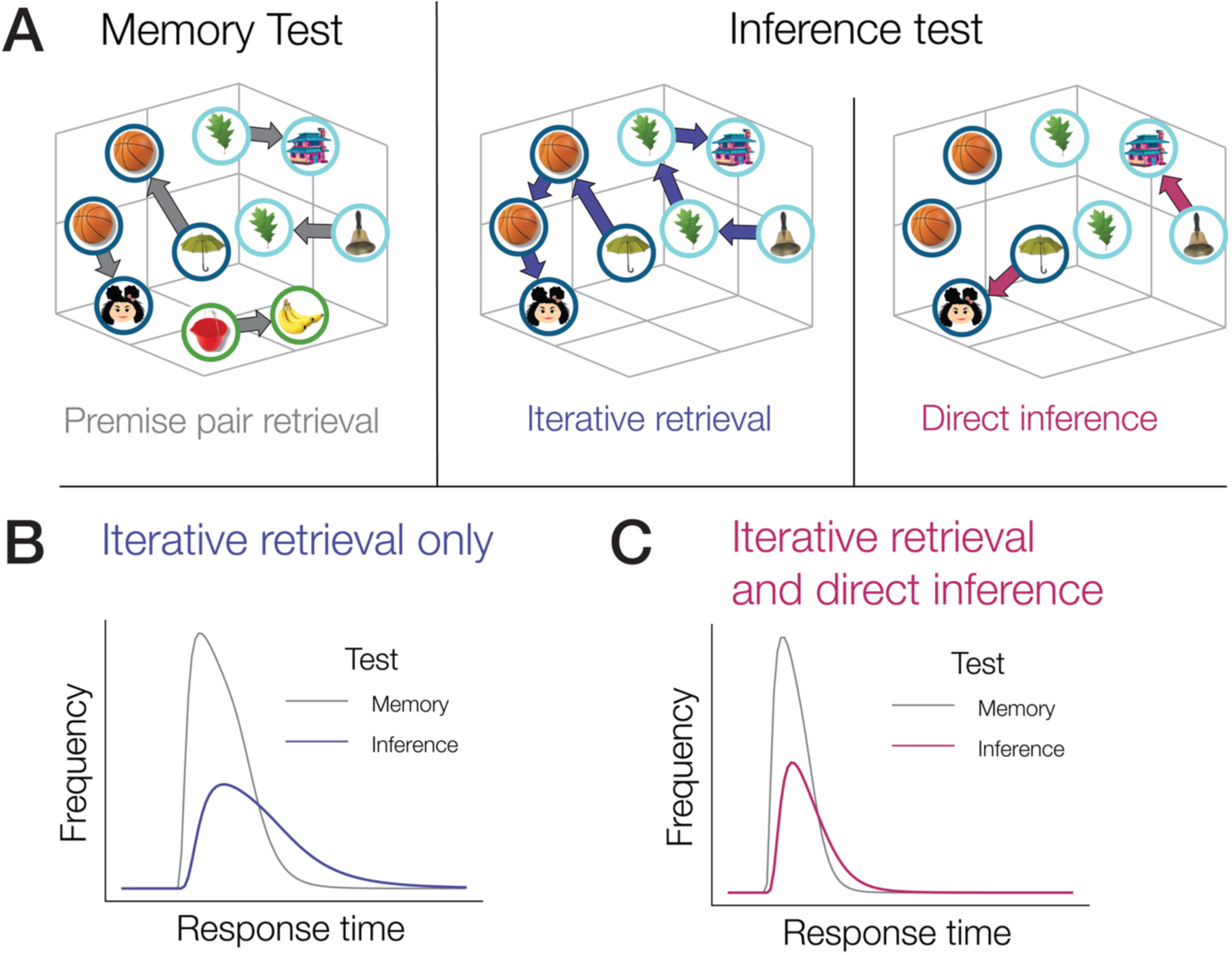
**(A)** Possible computations during the memory and inference tests. **(B)** If inference relies solely on iterative retrieval, response times should be slower for inference than memory. **(C)** If a direct route is also available, inference and memory should show more similar response times.

We modeled trial-level response times as arising from competition between accumulators corresponding to these processes. Critically, the drift rate (i.e., speed of evidence accumulation) for each process was modeled as a function of neural activation on each trial. Trial-wise activation from hippocampus and angular gyrus were entered as predictors of drift rate, allowing us to test whether these regions tracked distinct inference computations. To assess developmental change, we allowed both drift rates and neural–drift relationships to vary as a function of age. This approach enabled us to determine whether hippocampus and angular gyrus differentially supported variable-speed (consistent with iterative retrieval) and fixed-speed (consistent with direct inference) processes across development.

For exploratory analyses, we additionally included neural signals from ventral lateral prefrontal cortex, precuneus, intraparietal sulcus/superior parietal lobe, and parahippocampal cortex, given their involvement in inference and geometrically aligned representations (Morton et al., 2020). Results for these regions are reported in the **Supplement**. We estimated model parameters using hierarchical Bayesian inference and evaluated model fit using posterior predictive checks (see Morton (2026) for code and **Supplement** for full model specification, priors, and validation).

##### 2.9.3.1. Partial data exclusions

We excluded runs with excessive motion (more than one-third of volumes with mean framewise displacement > .30 mm) or incomplete acquisition due to scanner constraints or technical issues. Despite these exclusions, we retained all 74 participants. The hierarchical Bayesian framework accommodates missing data, allowing parameter estimates to leverage all available observations (Shiffrin et al., 2008). For trials with missing neural data, we imputed values by sampling from a standard normal distribution, ensuring that missingness did not bias model estimation.

##### 2.9.3.2. Model definition

The LBA model simulated responses across memory (BC, XY) and inference (AC) trials. On each trial, evidence accumulated toward competing response options, with separate accumulators capturing variable-speed (consistent with iterative retrieval) and fixed-speed (consistent with direct inference) processes (see **Supplement** for full specification). There were four accumulators in total: two accumulators for correct choices made with variable- or fixed-speed processes, *v*_1_ and *v*_2_, and two accumulators for incorrect choices, corresponding to lure items, with a drift rate of *v*_3_ for memory trials and *v*_4_ for inference trials.

We allowed drift rates for these processes to vary as a function of neural activation. Trial-level activation from hippocampus, angular gyrus, and exploratory regions predicted drift rate on each trial. We modeled these neural–drift relationships as linear functions of mean-centered age, allowing both the strength and direction of brain– behavior coupling to change across development. We applied informative priors to neural slope parameters to regularize estimates toward zero and reduce false-positive effects. We estimated model parameters using No U-Turn Sampling (NUTS; Hoffman & Gelman, 2011). Model performance was evaluated using posterior predictive checks, comparing simulated responses and response times to observed data (see **Supplement**).

## 3. Results

### 3.1. Initial associations provide stable premises for inference across development

Successful inference depends on the ability to form and retain the initial (AB) associations, which serve as premises for later reasoning judgments. We therefore tested whether memory for these pairs remained high across age and over time, from the time of the initial learning phase (which occurred pre-scan) to the final test phase (which occurred post-scan). The main effects model (accuracy ∼ age + test + (1 | participant)) best fit the data (conditional R^2^ = 0.46; marginal R^2^ = 0.10). Both age (χ^2^(1) = 8.69, *p* = .003) and test phase (χ^2^(1) = 6.50, *p* = .011) significantly predicted performance (**Fig. 4A**).

**Figure 4.**
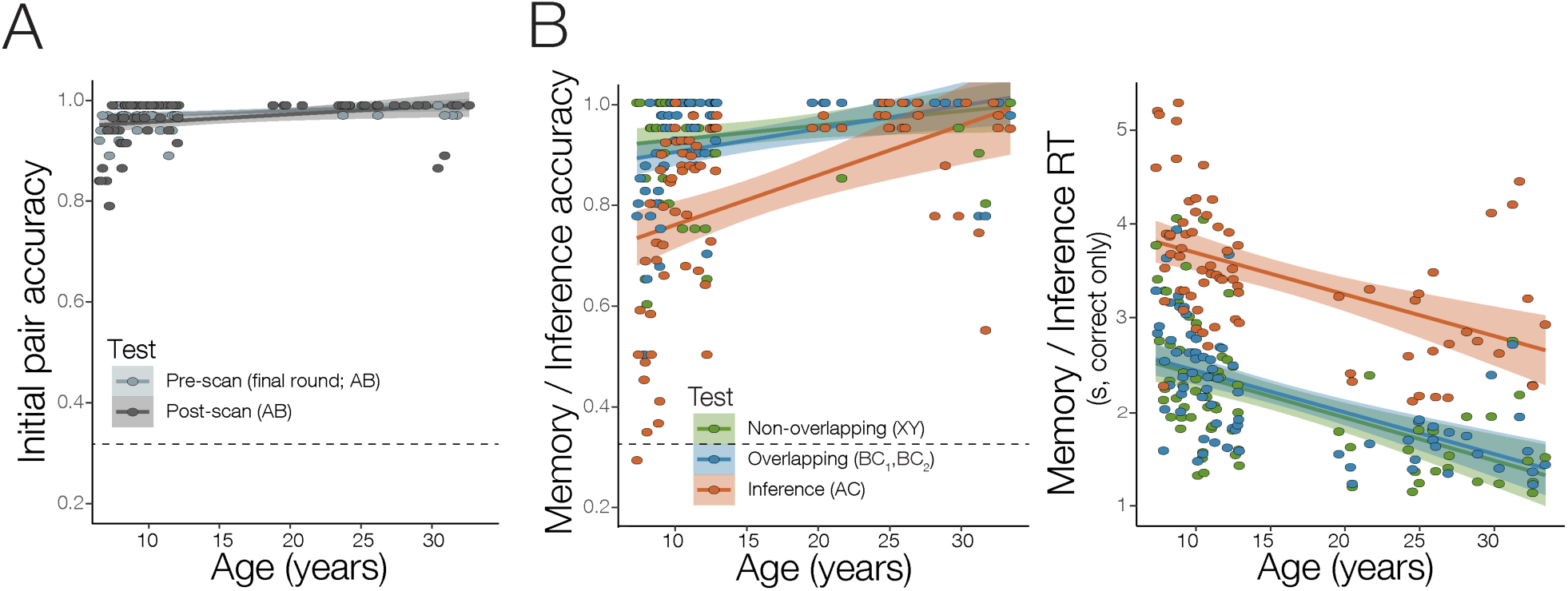
Developmental differences in inference exceed those for memory. **(A)** Accuracy for initial pairs (AB) remained near ceiling across age during both pre-scan learning and the post-scan test, indicating that participants formed and maintained the premise associations required for later inference. **(B)** Performance on non-overlapping memory (XY), overlapping memory (BC), and inference (AC) tests as a function of age. Accuracy was lower for inference than memory, particularly in children, indicating greater developmental differences for deriving relationships across associations than for retrieving directly learned pairs. Response times (correct trials only) decreased with age across all conditions but remained slower for inference than memory, even when accuracy approached ceiling in adults, consistent with additional computations supporting inference. Raw data are shown; dashed lines indicate chance performance.

Participants showed a small decline in performance from initial learning to the final test, consistent with modest forgetting, and performance increased slightly with age. Critically, however, initial pair memory remained near ceiling across groups and test phases (children: initial learning phase *M* = 98.04%, SEM = 0.36%, final test phase *M* = 96.60%, SEM = 0.71%; adults: initial learning phase *M* = 99.58%, SEM = 0.17%; final test phase *M* = 99.06%, SEM = 0.65%), indicating that participants formed and maintained robust memories of initial pairs. This result informs the interpretation of subsequent findings: developmental differences in inference cannot be attributed to failures in learning or maintaining the initial associations, but instead reflect differences in how these representations are used to derive inferences about novel relationships.

### 3.2. Inference shows a disproportionate developmental improvement relative to memory

Inference requires integrating information across separate experiences to arrive at a previously unobserved conclusion, whereas memory for directly learned associations involves retrieving a single directly experienced memory. Because inference depends on coordinating information across multiple representations, it should be more sensitive to developmental differences than retrieval of directly learned associations. To test this prediction, we compared performance across non-overlapping memory (XY), overlapping memory (BC), and inference (AC) tests.

Accuracy was best fit by an interaction model (accuracy ∼ age + test type + age × test type (1 | participant); conditional R^2^ = 0.69; marginal R^2^ = 0.29). Results showed a significant age x test type interaction (χ^2^(2) = 20.37, *p* ≤ .001), along with main effects of age (χ^2^(1) = 15.86, *p* < .001) and test type (χ^2^(2) = 110.24, *p* < .001; **Fig. 4B**). Marginal contrasts confirmed lower performance for inference compared to both overlapping and non-overlapping memory tests across all participants, (*ts*(214) ≤ −8.51, *ps* < .001), and no difference in performance between the memory tests (*t*(214) = 1.09, *p* = .283). The slope for age was also larger for inference than for both memory tests (*ts*(144) ≥ 3.25, *ps* ≤ .004), for which the slopes did not differ, *t*(144) = 1.09, *p* = .525. Supplemental analyses additionally showed a greater performance difference between the inference and memory tests in children (*ts*(214) ≤ −8.63, *ps* < .01) compared to adults (*ts*(214) ≤ −2.19, *ps* ≤ .03; see **Supplement**). Consistent with our prediction, participants were less accurate on inference than memory for directly learned pairs, and this difference was largest in children, indicating a disproportionate difficulty in deriving relationships across associations earlier in development. Although inference performance remained lower than memory across age, it approached ceiling in adults (*M* = 92.45%, SEM = 2.23%), indicating that the behavioral gap between inference and memory narrows across development. However, this ceiling-level performance in adults limits our ability to evaluate the mechanisms supporting successful inference using accuracy alone. To address this challenge, subsequent analyses focused on response time for correct trials. The substantial variability in this variable across all ages made it a more sensitive measure of the processes supporting performance.

Response times for correct trials were best fit by a main effects model (response time ∼ age + test type + (1 | participant); conditional R² = 0.85; marginal R² = 0.58). Both age (χ^2^(1) = 37.62, *p* < .001) and test type (χ^2^(2) = 629.39, *p* < .001) significantly predicted response times (**Fig. 4B**). While responses became faster with age across all conditions, marginal contrasts showed reliably slower response times for inference than both memory tests *ts*(216) ≥ 21.28, *ps* < .001), for which response times did not differ (*t*(216) = 0.87, *p* = .383). Critically, this response time cost persisted even when inference accuracy approached ceiling in adults (adult response times for inference test: *M* = 2.89 s, SEM = 0.14 s; overlapping memory test: *M* = 1.66 s, SEM = 0.07; and non-overlapping memory test: *M* = 1.62, SEM = 0.09), indicating that successful inference, on at least some trials, continues to require additional processing beyond retrieval of directly learned associations. This pattern suggests that response time provides a sensitive index of the computations supporting inference across development. Subsequent analyses collapse the non-overlapping (XY) and overlapping (BC) memory tests into a single memory condition, given similar response time performance across them and because our primary focus is on the additional computations required for inference versus direct memory across development.

### 3.3. Hippocampus supports inference across age, while parietal cortex tracks developmental differences

The observed differences in response times between inference and memory, suggest that inference relies on additional computations and that these computations may differ across development. To test this prediction, we next examined whether the neural systems supporting inference show dissociable patterns of engagement across age. Prior adult work suggests that inference can arise from at least two computations: an iterative retrieval process, supported through the retrieval of multiple associations (i.e., AB, BC), and a direct inference process, proposed to rely on a previously derived relationship (AC) among overlapping pairs (Schlichting & Preston, 2015; Morton et al., 2020).

We first identified regions that support inference independent of age by examining areas more strongly engaged during successful inference (AC) than memory (BC, XY), restricting analyses to correct trials to isolate processes supporting successful performance. This contrast revealed robust hippocampal activation that survived small-volume correction, along with distributed cortical activation including frontal and temporal regions associated with cognitive control and memory retrieval (**Fig. 5A**; see **Table S1** for a list of all statistically significant clusters) (Barredo et al., 2015; Friedman & Robbins, 2022; Marek & Dosenbach, 2019). These findings indicate that the hippocampus is consistently recruited when participants infer relationships across associations, regardless of age, consistent with a role in linking experiences through either iterative or direct retrieval.

**Figure 5.**
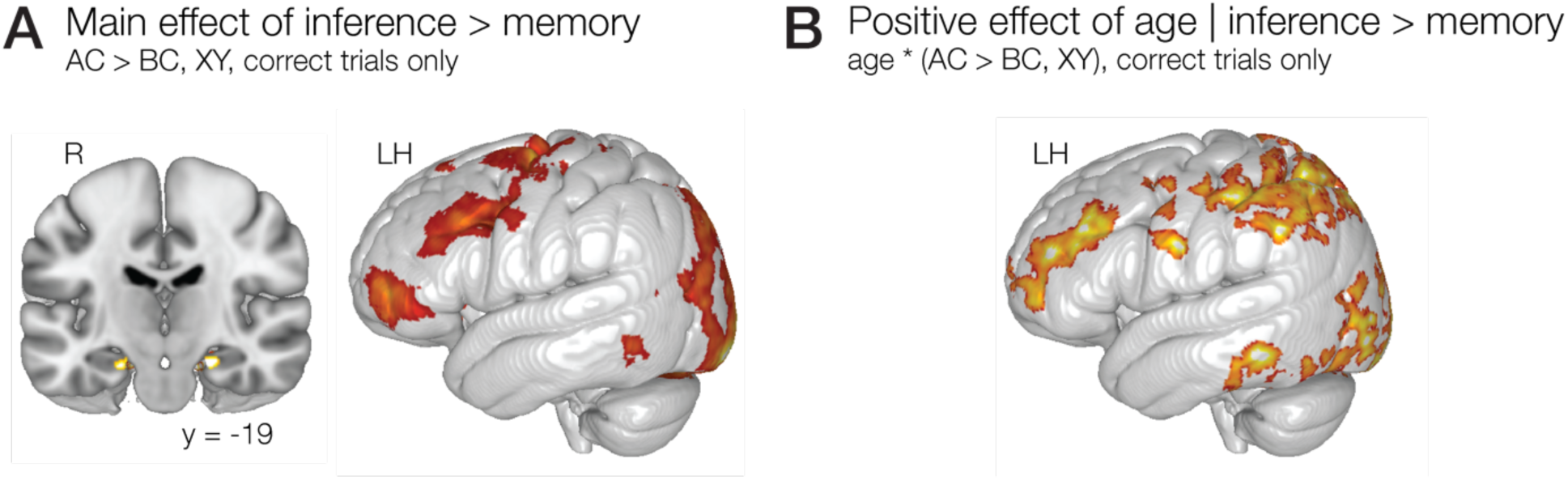
Distinct neural systems support inference across development and track developmental change. **(A)** Regions more strongly recruited during successful inference (AC) than memory (BC, XY) across all participants revealed robust activation in the hippocampus, along with distributed frontal and temporal regions. This pattern indicates that the hippocampus supports inference independent of age. **(B)** Regions showing greater recruitment for successful inference relative to memory (AC > BC, XY) with increasing age were observed in posterior parietal cortex, including angular gyrus, indicating an age-related increase in the engagement of parietal mechanisms during inference. Maps include clusters that survived correction for multiple comparisons across the whole brain, as well as those surviving small-volume correction within an *a priori* hippocampal ROI (*p* < .01; *a* < .05).

We next tested for regions showing an age-related increase in engagement for inference (AC) relative to memory (BC, XY), which would be consistent with a developmental shift in the computations supporting inference. This analysis revealed clusters in posterior parietal cortex that exhibited greater inference-related recruitment with increasing age (**Fig. 5B**; see **Table S1** for a list of all statistically significant clusters). This pattern aligns with the proposal that posterior parietal regions, including angular gyrus, support a more efficient, direct route to inference that becomes increasingly available across development. Together, these findings reveal a dissociation between the neural systems that support inference across age and those that track developmental differences in inference performance. The hippocampus appears to provide a stable mechanism for linking experiences, whereas posterior parietal cortex shows increasing engagement with age, consistent with a developmental shift toward more efficient inference computations.

### 3.4. Angular gyrus engagement indexes increasingly efficient inference with age

Whole-brain, univariate analyses indicated that posterior parietal cortex shows increasing engagement for successful inference relative to memory with age. We next asked whether this region—specifically the angular gyrus—supports a more efficient route to inference, in which relationships can be directly derived, rather than through the iterative retrieval of intermediate associations. If so, angular gyrus engagement should be selectively recruited during successful inference in adults and should be associated with faster correct inference responses.

To test these predictions, we examined whether angular gyrus activation varied by test type (memory, inference) and age. Angular gyrus activation was best fit by an interaction model (PE ∼ age + test type + age × test type (1 | participant); marginal R² = 0.06). A conditional R^2^ is not reported and the marginal R^2^ does not account for random effects because data were standardized within participant prior to analysis. This model yielded a significant age × test type interaction, χ^2^(1) = 6.04, *p* =.014 (**Fig. 6A**; main effect of test type: χ^2^(1) = 0.00, *p* = .979; main effect of age: χ^2^(1) = 2.41, *p* = .120). The interaction was driven by a significant slope for age during the inference (*t*(120) = 2.84, *p* = .005) but not memory tests (*t*(120) = −0.64, *p* = .524), indicating age-related differences in angular gyrus engagement for inference only. Children showed reduced engagement during inference (*M* = −18.88, SEM = 10.64), whereas adults showed robust engagement (*M* = 32.88, SEM = 13.79). In contrast, neither group reliably engaged angular gyrus during memory (children: *M* = 3.25, SEM = 6.53; adults: *M* = −9.13, SEM = 7.17)

**Figure 6.**
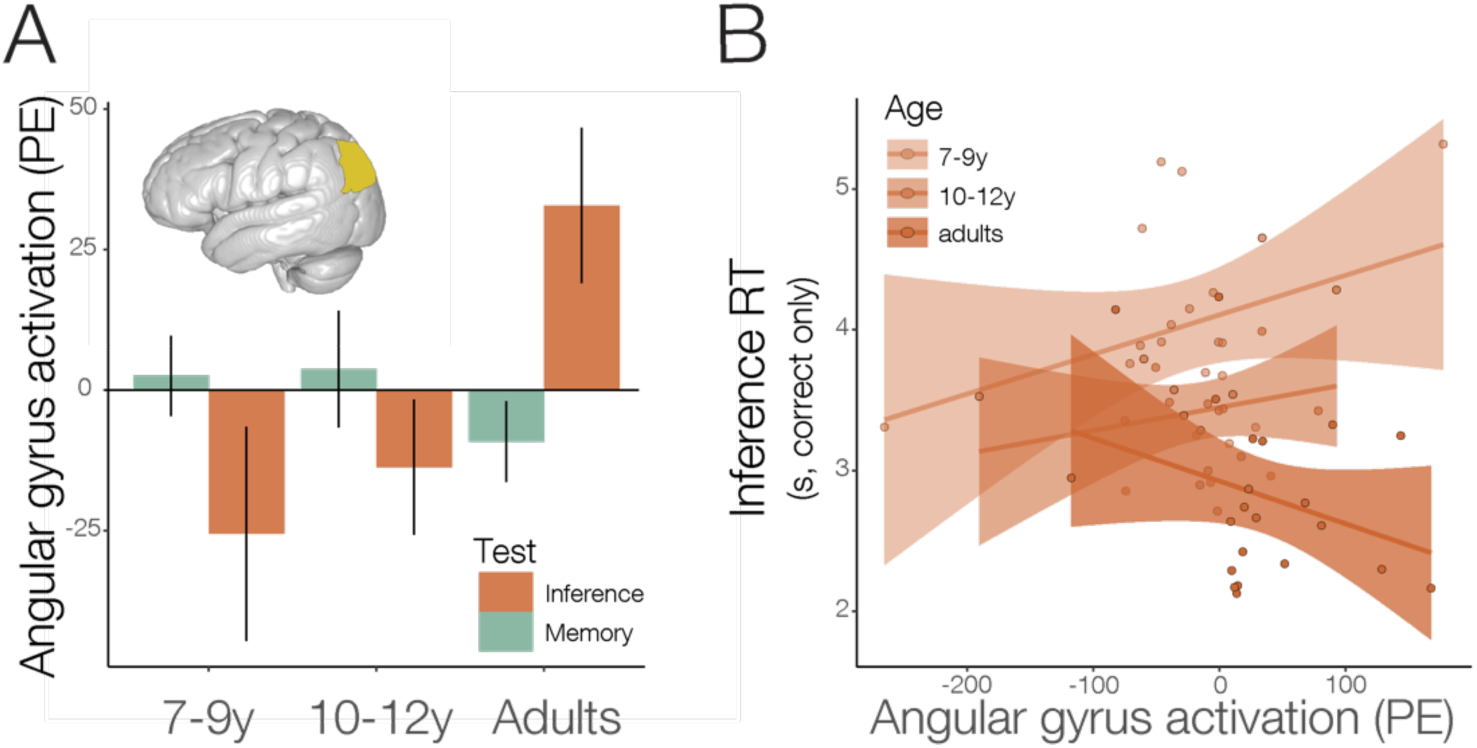
Angular gyrus engagement during inference increases with age and supports more efficient performance in adulthood. Age is treated continuously in all analyses but is binned here for visualization. Raw data are shown. **(A)** Mean activation and standard error (SE) within the angular gyrus ROI during correct memory (BC, XY) and inference (AC) trials for each age bin. Angular gyrus engagement increased with age selectively during inference, with adults showing robust recruitment and children showing no or reduced engagement. In contrast, angular gyrus was not reliably engaged during memory in any group. **(B)** Relationship between angular gyrus activation during inference and response time for correct inference trials. In adults, greater angular gyrus activation was associated with faster responses, whereas this relationship was absent or reversed in children, indicating that angular gyrus engagement supports more efficient inference with age.

We next tested whether angular gyrus engagement was associated with the efficiency of inference, indexed by response times for correct trials. Response times were best fit by an interaction model (response time ∼ age + angular gyrus PE + age × angular gyrus PE), which explained substantial variance (adjusted R² = 0.34, *F*(3,59) = 11.70, *p* < .001). This model revealed a significant age x angular gyrus PE interaction, *F*(1) = 6.18, *p* = .016 (**Fig. 6B**; main effect of age: *F*(1) = 24.58, *p* < .001; main effect of angular gyrus PE: *F*(1) = 0.05, *p* = .816). In adults, greater angular gyrus engagement tended to predict faster correct inference, consistent with a role in efficient, direct inference. This pattern was absent in children, for whom angular gyrus engagement was weakly associated with slower responses. Together, these findings suggest that angular gyrus contributes to a more efficient route to inference that emerges across development. While the hippocampus supports inference across age, angular gyrus engagement may index a distinct computation that enables faster, more efficient inference in adulthood.

### 3.5. Model-based analyses identify a developmental shift in the computations supporting inference

Behavioral and univariate results suggested that inference relies on at least two distinct computations: a slower, iterative retrieval process supported by the hippocampus and a faster process that becomes increasingly associated with angular gyrus engagement in adulthood. In the context of the present task, these computations correspond to retrieving premise pairs (AB, BC) sequentially to derive the AC relationship, versus accessing the inferred (AC) relationship more directly. To directly test this hypothesis, we used a multilevel decision model to dissociate these computations and examine whether trial-by-trial variability in hippocampal and angular gyrus activation tracked different routes to successful inference across development (**Fig. 7A**). The model formalized iterative retrieval as a variable-speed process, which is slower for inference than memory, and direct inference as a fixed-speed process, which operates similarly across conditions (**Fig. 3**). Critically, this framework allowed us to test whether neural activity predicted the speed of these distinct processes on a trial-by-trial basis and whether these relationships changed with age. We calculated 94% high-density intervals (HDIs) to quantify the uncertainty surrounding estimated effects.

**Figure 7.**
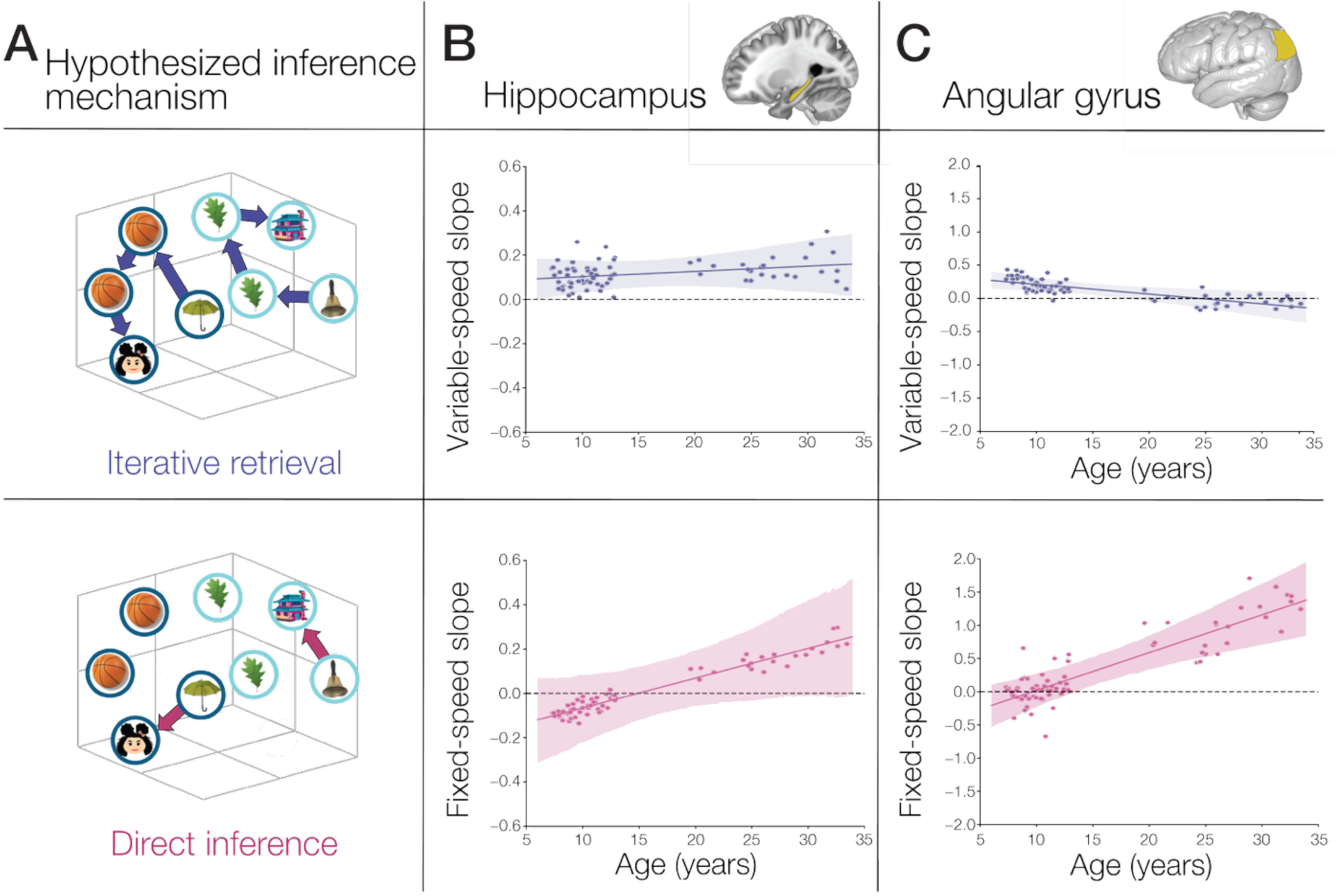
Model-based analyses reveal a developmental shift in the computations supporting inference. **(A)** A multilevel decision model was used to dissociate the two hypothesized inference computations: iterative retrieval which required retrieving multiple associations (AB, BC), and direct inference which required retrieving one association (AC). Iterative retrieval was therefore formalized as a variable-speed process (producing slower responses for inference than memory) and direct inference as a fixed-speed process (producing similar response times across conditions). **(B)** Hippocampal activation predicted the speed of the variable-speed process, a relationship that was stable across age (top panel). Hippocampal activation also became increasingly associated with the fixed-speed process with age (bottom panel). **(C)** Angular gyrus activation tracked the variable-speed and fixed-processes differently with age. Its association with the variable-speed process decreased with age (top panel), while its association with the fixed-speed process increased with age (bottom panel). Furthermore, in adults, greater angular gyrus activation predicted faster operation of the fixed-speed process, a relationship not observed in children. Together, these results suggest a developmental shift in the computations supporting inference. In younger participants, the hippocampus supported correct inference through iterative retrieval. In adults, more efficient inference through direct retrieval was associated with angular gyrus, with increasing contributions of the hippocampus.

Consistent with prior work (Morton et al., 2020), hippocampal activation tracked the variable-speed process associated with iterative retrieval (slope 94% HDI: [0.0574, 0.1741]; **Fig. 7B**, **top panel**). This relationship did not vary reliably with age (slope change 94% HDI: [−0.0044, 0.0094]), indicating that the hippocampus supports iterative retrieval similarly in children and adults. Hippocampal activation did not reliably predict the fixed-speed process associated with more direct AC retrieval (slope 94% HDI: [−0.1093, 0.1271]), and this relationship did not vary reliably with age. However, parameter estimates suggested a weak positive age-related trend (slope change 94% HDI: [−0.0006, 0.0270]; **Fig. 7B**, **bottom panel**). Although this effect was not reliable and should be interpreted with caution, the direction of this trend raises the possibility that hippocampal contributions to inference may become less strictly tied to iterative retrieval with age.

To further investigate this possibility, we directly compared the relationship between hippocampal activation and the fixed-speed process in children and adults. Specifically, we compared the model-direct slope parameters, which quantify the strength of the relationship between trial-level neural activation and the speed of the fixed-speed process. For each sample of the posterior distribution, we calculated the mean slope for children and adults and then computed the difference between groups. This analysis revealed a stronger positive relationship between hippocampal activation and the fixed-speed process in adults versus children (94% HDI of slope difference: 0.0038 to 0.4511). These results thus indicate that hippocampus supports an iterative retrieval mechanism that remains stable across development, while also becoming increasingly associated with a direct inference process in adulthood.

Furthermore, angular gyrus activation tracked both the variable- and fixed-speed processes in a developmentally specific manner. Angular gyrus activation predicted the speed of the variable-speed (iterative) process overall (slope 94% HDI: [0.0391, 0.2161]), but this relationship decreased with age (slope change 94% HDI: [−0.0259, −0.0032]; **Fig. 7C**, **top panel**), indicating that angular gyrus contributes to iterative retrieval in children but not adults. Notably, angular gyrus activation also predicted the speed of the fixed-speed process associated with direct inference (slope 94% HDI: [0.1305, 0.5649]), and this relationship increased with age (slope change 94% HDI: [0.0305, 0.0837]; **Fig. 7C**, **bottom panel**). In adults, greater angular gyrus engagement was associated with faster operation of this process, whereas this relationship was absent in children.

We further tested these developmental differences by directly comparing the neural slope parameters between children and adults, using the same approach described for the hippocampus (see **Table S2**). Consistent with the continuous-age analyses, the relationships between angular gyrus activation and the variable-speed process was stronger in children than adults (94% HDI of slope difference: [−0.4279, −0.0772]). In contrast, the relationship between angular gyrus activation and the fixed-speed process was stronger in adults than children (94% HDI of slope difference: [0.5664, 1.3886]). Together, these findings provide converging evidence for a developmental difference in the neural computations associated with angular gyrus activation. Whereas angular gyrus activation was more strongly associated with iterative retrieval in children, it was more strongly associated with direct inference in adults. These developmental differences may contribute to increasingly efficient inference with age, alongside potential contributions of the hippocampus to direct inference with age. See **Supplement** for results from exploratory analyses of neural signals from other regions.

## 4. Discussion

Drawing inferences from past experience is fundamental to learning. This ability allows individuals to extend beyond directly observed information to guide decisions, generalize knowledge, and support academic achievement (Esposito & Bauer, 2017; Schlichting & Preston, 2015; Varga et al., 2019). Despite its central role in cognition, the neurocomputational mechanisms that support inference—and how they differ across development—remain largely unknown. One account proposes that children and adults rely on qualitatively different routes to inference (Bauer et al., 2015, 2021; Schlichting et al., 2017; Shing et al., 2019; Varga & Bauer, 2013). In this framework, children iteratively retrieve related experiences, while adults can access representations that directly link related events, enabling a more efficient computation. Critically, this distinction has remained largely theoretical, as prior work has not directly tested the computations engaged at the actual time of inference. The present study addresses this gap using model-based neuroimaging to dissociate the neural computations supporting inference in children and adults. Across development, hippocampal engagement tracked an iterative retrieval mechanism and did not vary with age. In contrast, angular gyrus engagement increasingly tracked a faster, direct inference mechanism with age. There were also stronger associations between this direct inference mechanism and activation in angular gyrus and hippocampus in adults relative to children. These findings identify a possible shift in the neural computations supporting inference across development. More specifically, they indicate stable hippocampal contributions to iterative retrieval alongside age-related changes in parietal cortex that support more efficient inference.

These findings provide a mechanistic account of well-established developmental differences in inference performance. Consistent with prior work, children were less accurate and slower when making inference judgments, and their memory for overlapping events was less tightly coupled to inference performance (Abolghasem et al., 2023; Schlichting et al., 2017, 2022; Shing et al., 2019). Our neuroimaging results suggest that these behavioral differences reflect differences in the computations supporting inference, rather than differences in memory strength alone. In particular, children appear to rely on iteratively retrieving related experiences at the time of decision. In contrast, adults appear to additionally access representations that directly link related events, enabling more efficient inference. This interpretation helps reconcile prior behavioral and neuroimaging findings. Behavioral work suggests that children tend to construct inferences on demand, whereas adults are more likely to form direct links between events in advance (Bauer et al., 2015, 2021; Varga & Bauer, 2013). Converging evidence from neuroimaging work shows that adults often reactivate prior events during new learning, supporting the formation of integrated representations (Schlichting & Preston, 2015). By contrast, children show weaker or more transient reactivation and maintain more differentiated representations of related events (Schlichting et al., 2022; Varga et al., 2025). The present findings extend this work by demonstrating that these representational differences parallel distinct computations at the time of inference.

The present findings also help clarify the role of posterior parietal cortex in reasoning. Prior work has linked this region to the development of reasoning abilities (Crone et al., 2009; Wendelken et al., 2011, 2016) across a range of tasks in adults, including analogical (Hobeika et al., 2016), arithmetic (Ischebeck et al., 2009), and spatial (Sack, 2009) reasoning. However, these accounts have primarily characterized when posterior parietal cortex is engaged across development, rather than what computations it supports. Our findings address this gap by identifying a specific computational role for angular gyrus in supporting more efficient inference with age. In the context of memory-based reasoning, angular gyrus engagement tracked a faster, direct inference mechanism that became increasingly prominent with age. This pattern suggests that developmental changes in parietal cortex may support a broader shift toward more efficient reasoning strategies across domains. An important question for future work is whether this same computational mechanism contributes to age-related improvements in non-memory forms of reasoning. Establishing such a link would provide a more unified account of reasoning development and clarify how domain-general and memory-specific processes interact to support flexible cognition.

Posterior parietal cortex has also been widely implicated in memory retrieval in adults, though its precise role in retrieval operations remains debated (e.g., Cabeza et al., 2008; Olson & Berryhill, 2009; Shimamura, 2011; Wagner et al., 2005). One influential account proposes that this region supports the temporary maintenance of retrieved information, buffering memories online to enable further evaluation and decision-making (Baddeley, 2000; Vilberg & Rugg, 2008). Under this framework, inference would be expected to rely on angular gyrus maintaining and combining multiple retrieved associations, consistent with an iterative retrieval process.

Our findings are not consistent with this account in adulthood. Rather than tracking a variable-speed process indicative of iterative retrieval, angular gyrus activation tracked a fixed-speed process, consistent with direct inference from representations that link related events (Morton et al., 2020). Access to these representations may reduce the need to temporarily maintain and combine individual associations in working memory, as would be required for iterative retrieval. These findings refine current accounts of parietal contributions to memory by suggesting that angular gyrus function depends on how related experiences are represented and the computations required to access them. When memories are represented separately, angular gyrus may contribute to buffering and combining retrieved associations. In contrast, direct access to integrated or geometrically aligned representations may support more efficient inference without these intermediate operations. Together, our findings suggest that developmental changes in angular gyrus function may reflect a shift in the computations associated with memory retrieval, rather than a uniform increase in retrieval-related processing.

Our findings suggest that angular gyrus supports direct inference in adults, enabling access to representations that directly link related events. These representations may arise through at least two mechanisms. One possibility is that inference relies on integrated representations formed during encoding, in which related events are combined into an overlapping memory trace (**Fig. 1A**, middle panel) (Schlichting & Preston, 2015, 2016; Zeithamova et al., 2012; Zeithamova & Preston, 2010). A second possibility is that inference is supported by navigating structured representations, in which related events are geometrically aligned based on shared features or relationships (**Fig. 1A**, right panel). In this case, inference can proceed efficiently by traversing these aligned representations without retrieving each association separately. Consistent with this latter account, prior work has shown that adults form geometrically aligned representations in posterior parietal cortex that support successful inference (Morton et al., 2020).

A key implication of the present findings is that children may rely on a different representational format to support inference. Rather than accessing integrated or aligned representations, children may depend on iteratively retrieving and combining individual associations at the time of decision. Our computational response time model suggests that angular gyrus is associated with iterative retrieval in children, potentially reflecting a role in buffering associations online (Baddeley, 2000; Vilberg & Rugg, 2008). However, our univariate analysis showed reduced angular gyrus activation during successful inference relative to premise pair memory in children. Although these findings may initially seem inconsistent, they reflect different aspects of angular gyrus function. The univariate analyses assessed overall differences in activation between inference and memory, whereas the computational model examined how trial-by-trial variability in activation related to the speed of distinct inference mechanisms. Thus, reduced overall angular gyrus activation during inference does not preclude a positive relationship between activation and the speed of iterative retrieval. For example, angular gyrus activation may be reduced overall during inference, but decrease less—or even increase—on trials when greater activation is associated with faster operation of the variable-speed process.

Importantly, these findings clarify how angular gyrus engagement relates to inference performance in children. Although the univariate analysis revealed less inference-related recruitment in children than adults, with reduced engagement during inference relative to memory in the youngest children, these findings do not imply that angular gyrus is uninvolved in children’s inference. In our computational model, trial-level angular gyrus activation was positively associated with the variable-speed process in children, consistent with iterative retrieval. That children engaged angular gyrus when they were also slower to make correct inference decisions is also consistent with this mechanistic interpretation. In contrast, greater angular gyrus activation in adults was associated with faster inference responses and the fixed-speed process. These opposing patterns suggest developmental differences in angular gyrus function, although they do not establish that angular gyrus directly implements iterative retrieval in children. Such differences are consistent with the region’s protracted development. Angular gyrus peaks in volume between late childhood and early adolescence (Gogtay et al., 2004; Seghier, 2012; Westlye et al., 2010) and its connectivity with the hippocampus also continues to develop into emerging adulthood (Lebel & Beaulieu, 2011). These changes may support the formation and use of integrated or geometrically aligned representations with age, enabling a more efficient, direct route to inference.

Although developmental changes in inference were primarily associated with angular gyrus, the hippocampus provided a stable computational foundation for inference across age. Hippocampal engagement tracked a variable-speed process consistent with iterative retrieval in both children and adults. This pattern suggests that the hippocampus supports inference by retrieving and combining individual experiences at the time of decision, a mechanism that remains available across development. Given its well-established role in memory retrieval (e.g., Eichenbaum et al., 2007; Squire et al., 2007; Wiltgen et al., 2010), this contribution is consistent with accounts that emphasize the hippocampus as supporting flexible access to relational information. From a developmental perspective, this mechanism may play different roles across age. In childhood, when parietal representations are not yet available to support direct inference, iterative retrieval may provide a primary route to successful inference. In adulthood, this mechanism may still be maintained and recruited when more direct representational routes are unavailable or fail. Hippocampal engagement may therefore support a flexible, albeit slower, pathway that can be used across contexts, even as other systems enable more efficient computations with development.

Model estimates also provided limited evidence that hippocampal engagement may also relate to a direct inference mechanism in adulthood. Although this effect should be interpreted with caution, it suggests that hippocampal contributions may be less exclusively tied to an iterative retrieval mechanism with age. This interpretation is broadly consistent with evidence for continued development of hippocampal function and its role in supporting increasingly efficient memory retrieval (DeMaster et al., 2016; Ghetti & Bunge, 2012; Sastre et al., 2016), as well as with findings linking hippocampal structure to inference performance across development (Schlichting et al., 2017). Importantly, this pattern was only evident in the model-based analyses and was not observed in univariate activation, underscoring the value of approaches that can distinguish between computational processes. Future work that combines model-based approaches with measures of neural representation at the time of inference will be important for clarifying whether, and under what conditions, the hippocampus contributes to a direct inference mechanism.

While the present work extends our understanding of the neural computations supporting inference development, important questions remain. We examined developmental differences in both the recruitment of hippocampus and angular gyrus during inference as well as the relationship between their activation and distinct computational processes. Our findings show that angular gyrus engagement both increases with age and becomes more strongly associated with the more efficient, direct inference mechanism, supporting a developmental shift in how this region contributes to inference. However, our approach cannot determine how frequently individuals engage in direct inference versus iterative retrieval on a trial-by-trial basis. For example, individuals may rely on iterative retrieval of overlapping associations for triads wherein the premises (AB, BC) are represented separately, but access inferred relationships more directly when premise associations are integrated or geometrically aligned. Testing this possibility will require measuring the neural representations of individual triads before and after learning (Schlichting et al., 2015; Varga et al., 2025), allowing future studies to relate representational structure for individual triads to computations engaged during triad-specific inference decisions (Morton et al., 2020). Such work will clarify how developmental changes in memory representation influence the flexible use of different inference mechanisms.

Beyond hippocampal and angular gyrus contributions, we also observed age-related increases in the recruitment of a frontoparietal network during successful inference relative to memory (see **Table S1**). Although not the primary focus of the present study, this pattern is consistent with a role for cognitive control processes in supporting the execution of inference. Frontoparietal regions are known to support strategic retrieval operations, including the elaboration of retrieval cues, the selection of relevant information, and the monitoring of retrieved content (Badre et al., 2005; Badre & Wagner, 2007; Barredo et al., 2015; Cabeza et al., 2008; Rugg, 2022; Wagner et al., 2005). In the context of the present findings, these processes may help coordinate the retrieval and combination of information required for inference, particularly when multiple associations must be accessed or evaluated. From a developmental perspective, increased recruitment of frontoparietal regions may reflect improvements in the ability to efficiently engage and regulate these control processes during inference. This interpretation aligns with evidence for continued maturation of frontoparietal networks through adolescence (Grayson & Fair, 2017; Gu et al., 2015; Marek & Dosenbach, 2019). Notably, these regions are unlikely to implement the core computations underlying inference themselves. Instead, they may support the selection and deployment of hippocampal and angular gyrus mechanisms, enabling more efficient use of available representations with development.

Developmental improvements in inference have long been attributed to changes in how related events are represented and accessed in memory, yet this account has remained largely untested at the level of underlying computation. By combining model-based approaches with neuroimaging, an approach that remains rare in developmental research, the present study interrogated these mechanisms at the time of inference. This framework suggests that improvements in inference do not reflect a uniform strengthening of a single process, but rather a shift in the neural computations that support behavior. Children engaged an iterative retrieval mechanism supported by the hippocampus, while adults additionally engaged a more efficient, direct inference mechanism supported by angular gyrus and possibly hippocampus. These findings revise current accounts of memory development by showing that inference is supported by multiple neural pathways that differ in efficiency and emerge along distinct developmental trajectories. They also highlight a previously underappreciated role for posterior parietal cortex in memory-based reasoning. The development of this region may not only reflect stronger memory, but a reorganization of how memory is used to guide behavior. More broadly, this work demonstrates the value of integrating computational models with neuroimaging to uncover mechanisms not observable from behavior or neural activation alone. Given the central role of inference in learning—and its links to academic success—these findings provide a foundation for understanding how knowledge is built from experience and how these processes might be supported across development.

## 5. Declaration of generative AI and AI-assisted technologies in the manuscript preparation process

During the preparation of this work C. Coughlin used Google AI and ChatGPT as general search engines and to provide guidance with analyses and data visualization, as well as assistance with related code. ChatGPT was also used minimally to improve language and readability (e.g., phrasing suggestions or improvements, clarity/organization assistance, grammar checks). DALL-E was used to generate face and scene stimuli images for Fig. 2 and the graphical abstract, as those used in the actual experiment are under copyright protection. After using these tools, C. Coughlin reviewed or edited content as needed and takes full responsibility for the content of the published article.

## 6. Acknowledgements

We thank the many children, adults, and families who generously gave their time to support developmental science and make this research possible. We are also grateful to the members of the Preston Lab who provided valuable insights and assistance, especially Nicole Varga, Robert Molitor, and many undergraduate research assistants. Finaly, we appreciate the statistical guidance provided by Drs. Alexander Demos and Ryne Estabrook.

## Data and code availability

Data and analysis code for all but the GLM univariate analysis are available at https://osf.io/hku4q/overview?view_only=519523a5eca742c386a1d370c2f7267b and https://doi.org/10.5281/zenodo.19364577.

