## Supplementary Materials for "Children and adults use distinct neurocognitive mechanisms to support successful memory-based inference"

#### Supplementary Methods

##### *Participants*

A total of 121 participants enrolled in the multi-experiment study on day 1. Of these, 44 did not return for day 3, during which the present experiment was administered. Participants failed to return due to exclusion (n = 31), voluntary withdrawal (n = 8), or loss to follow-up (n = 5). Exclusion criteria included: below-average IQ (defined as scoring > 2 SD below the mean on the Wechsler Abbreviated Scale of Intelligence, Second Edition; Wechsler, 2011); psychological symptoms outside the typical range (children: scores in the clinical range on the Child Behavior Checklist; Achenbach, 1991; adults: scores > 1 SD above the normative mean on the Symptom Checklist 90-Revised; Derogatis, 1977); disclosure of a developmental or psychological disorder; non-native English speaker (learned after age 3); MRI contraindication; discomfort with the mock MRI environment; poor MRI data quality on day 2; prior participation in a related study; behavioral or technical issues during earlier sessions; or age outside remaining recruitment needs. The final sample for the present experiment included 77 participants. Exclusion information for this subset is reported in the main text.

##### *Experimental Procedures*

The present experiment was conducted on day 3 of a multi-experiment study and involved administration of the associative inference task (described in detail in the main text). Here, we briefly describe procedures across days 1–3 to provide context for the full study design. On day 1, participants were screened for IQ within the normal range (WASI-II; Wechsler, 2011) and for mental health symptoms below the clinical range (children: Child Behavior Checklist, Achenbach, 1991; adults: SCL-90-R, Derogatis, 1977). Participants also completed surveys assessing their mental health and background, were introduced to the MRI environment using a mock scanner, and completed a battery of cognitive tasks.

Cognitive tasks unrelated to the present experiment included the alternative uses task (Guilford et al., 1960) and the episodic specificity induction task (Madore et al., 2014). The alternative uses task assessed divergent thinking by asking participants to generate as many novel uses as possible for two everyday objects. The episodic specificity induction task assessed the development and relationship between remembering past events and imagining future events; participants remembered past events, imagined future events, and described pictures following either a control induction (math problems) or a memory retrieval induction.

Cognitive tasks related to the present experiment included the know game (see main text) and brief practice versions of the initial learning and non-overlapping learning phases of the associative inference task (using stimuli distinct from those used in the main task). This practice familiarized participants with the task structure (e.g., trial timing and response procedures) without revealing the presence of overlapping associations, which would occur during the full task on day 3.

Participants who remained eligible and were interested in completing the MRI portion of the study returned for day 2 approximately five weeks later (M = 4.81 weeks, SEM = 0.41). On day 2, participants completed anatomical scans, localizer functional MRI scans (0–3 runs), and resting-state functional MRI scans (0–2 runs). The localizer task was a 1-back paradigm involving images of faces, scenes, objects, and scrambled objects, none of which overlapped with stimuli used in the associative inference task. These localizer and resting-state data are not analyzed here and reported only to depict the full experimental procedure. Participants who continued to remain eligible and interested returned for day 3 approximately one week later (M = 0.99 weeks, SEM = 0.09). On this day, they completed the associative inference task, as described in the main text.

#### ***Formalization of Computational Model of Response Time***

To isolate the computations supporting performance on the memory and inference tests, we modeled responses as a competition between multiple decision accumulators. On each trial, the model response was determined by the accumulator that first reached a decision threshold, and response time was defined by the time required to reach that threshold. This framework allowed us to dissociate contributions of processes consistent with iterative retrieval and more direct inference mechanisms at the level of response dynamics.

Our modeling framework is based on the linear ballistic accumulator (LBA) model (Brown & Heathcote, 2008), which assumes that, on each trial, the starting point $k$ of each accumulator is drawn randomly from a uniform distribution on the interval $[0, A]$. Each accumulator then follows a line with a slope of $d$ until it reaches the response threshold $b$. On each trial, the slope $d$ of accumulator $i$ is drawn from a normal distribution with mean $v_i$ and standard deviation $s$ (here, fixed at 1). The time for an accumulator to reach the threshold is $(b - k)/d$.

As derived in the initial description of the LBA model (Brown & Heathcote, 2008), the probability density function (PDF) for accumulator $i$ at time $t$ is:

$$f_i(t) = \frac{1}{A}\left[-v_i\Phi\left(\frac{b-A-tv_i}{ts}\right) + s\phi\left(\frac{b-A-tv_i}{ts}\right) + v_i\Phi\left(\frac{b-tv_i}{ts}\right) - s\phi\left(\frac{b-tv_i}{ts}\right)\right]$$

Where $\phi$ and $\Phi$ are the probability density function and cumulative distribution functions, respectively, of the standard normal distribution. The cumulative distribution function (CDF) for accumulator $i$ at time $t$ is:

$$F_i(t) = 1 + \frac{b-A-tv_i}{A}\Phi\left(\frac{b-A-tv_i}{ts}\right) - \frac{b-tv_i}{A}\Phi\left(\frac{b-tv_i}{ts}\right) + \frac{ts}{A}\phi\left(\frac{b-A-tv_i}{ts}\right) - \frac{ts}{A}\phi\left(\frac{b-tv_i}{ts}\right)$$

The PDF for accumulator $i$ hitting the threshold first, at time $t$, is the probability of accumulator $i$ finishing at time $t$, conditional on the other accumulators not having finished yet:

$$\text{PDF}_i(t) = f_i(t) \prod_{j \neq i} \left(1 - F_{j(t)}\right)$$

Each three-alternative forced-choice (3AFC) test was modeled as a competition between four accumulators. Two accumulators, corresponding to variable-speed and fixed-speed processes, with average drift rates $v_1$ and $v_2$, respectively, supported the correct choice. Two additional accumulators represented the two incorrect choices, which corresponded to the lure items; these accumulators had an average drift rate of either $v_3$ (on memory test trials) or $v_4$ (on inference test trials).

The variable-speed accumulator is designed to simulate an iterative retrieval mechanism, which is assumed to be relatively fast for retrieving directly learned, premise associations (BC, XY) and slower for retrieving inferred associations (AC) (Morton et al., 2020). The variable-speed accumulator has an average drift rate $v_1$ for memory test trials (both BC and XY) and a slower drift rate $v_1 r$ for inference test trials (AC); $r$ is constrained to be between 0 and 1. The fixed-speed accumulator is designed to simulate a direct inference mechanism that may retrieve either direct or indirect associations with a fixed speed; this process may correspond to the retrieval of an integrated memory or geometrically-aligned representation (Morton et al., 2020). The fixed-speed accumulator has an average drift rate $v_2$. For both memory (BC, XY) and inference (AC) tests, non-decision time (including any time not related to the decision-making process, such as the time to perceive the test stimuli) was modeled as a fixed time interval $\tau$.

The probability of a correct response at time $t$ was:

$$P(\text{correct}, t) = \text{PDF}_1(t - \tau) + \text{PDF}_2(t - \tau)$$

The probability of an incorrect response at time $t$ was

$$P(\text{incorrect}, t) = 2\text{PDF}_3(t - \tau)$$

for memory tests and

$$P(\text{incorrect}, t) = 2\text{PDF}_4(t - \tau)$$

for inference tests.

The model was implemented in Python 3.10 using Morton (2026). We used Bayesian sampling to estimate parameters, using the No U-Turn Sampler (NUTS) implemented in PyMC 4.2.0. We fixed $s = 1$ to ensure parameter identifiability. The threshold parameter $b_i$ for each participant $i$ was modeled as a linear function of the mean-centered age of each participant in years, according to

$$b_i \sim \exp\big(\mu_b + \sigma_b \text{Normal}(0, 1)\big),$$

where:

$$\mu_b = b_{\beta 0} + b_{\beta 1}\text{age}$$
$$\sigma_b \sim \text{Cauchy}(0.5)$$

The $b_i$ parameters were defined in terms of log values to ensure that they would be strictly positive. Priors for age coefficients were:

$$b_{\beta 0} \sim \text{Normal}(2, 0.5)$$
$$b_{\beta 1} \sim \text{Normal}(0, 0.1)$$

The $\tau$, $A$, and $r$ parameters were fixed across subjects to ensure the stability of parameter estimates. Priors were:

$$\tau \sim \text{Normal}(0.5, 0.5) \epsilon [0, 1]$$
$$A \sim \text{Normal}(4, 2) \epsilon [0, \infty)$$
$$r \sim \text{Normal}(0.5, 0.5) \epsilon [0, 1]$$

Drift rates for the variable-speed and fixed-speed processes were free to vary from trial to trial based on neural signals measured during each test trial. The drift rate for subject $i$, trial $j$, and accumulator $k$, based on each neural signal $n$, was

$$v_{k,ij} = \gamma_{ik} + \sum_{n} \gamma_{n,ik} z_{n,ij},$$

where $z_{n,ij}$ is the neural signal for subject $i$, trial $j$, and signal $n$. Each signal was calculated as the first eigenvariate of activation within a given functional region of interest, which was then z-scored within run. The mean drift rate for each subject $i$ and accumulator $k$ was a linear function of age, according to

$$\gamma_{ik} \sim \mu_k + \sigma_k \text{Normal}(0, 1),$$

where:

$$\mu_k = \theta_{k,\beta 0} + \theta_{k,\beta 1}\text{age}$$
$$\sigma_k \sim \text{Cauchy}(0.5)$$

Priors for age coefficients were:

$$\theta_{k,\beta 0} \sim \text{Normal}(3, 1)$$
$$\theta_{k,\beta 1} \sim \text{Normal}(0, 0.1)$$

The slope parameter for each subject $i$ relating neural signal $n$ to the drift rate of accumulator $k$ on individual trials followed

$$\gamma_{n,ik} \sim \mu_{kn} + \sigma_{kn} \text{Normal}(0, 1),$$

where:

$$\mu_{kn} = \theta_{k,n,\beta 0} + \theta_{k,n,\beta 1}\text{age}$$
$$\sigma_{kn} \sim \text{Cauchy}(0.5)$$

Priors for age coefficients were:

$$\theta_{k,n,\beta 0} \sim \text{Normal}(0, 1)$$
$$\theta_{k,n,\beta 1} \sim \text{Normal}(0, 0.1)$$

The incorrect accumulators had drift rates that were free to vary depending on whether the test was memory or inference. The drift rate for subject $i$, condition $c$ was

$$v_{ic} \sim \mu_c + \sigma_c \text{Normal}(0, 1),$$

where:

$$\mu_c = \beta_{c0} + \beta_{c1}\text{age}$$
$$\sigma_c \sim \text{Cauchy}(0.5)$$

Priors for age coefficients were:

$$\beta_{c0} \sim \text{Normal}(3, 1)$$
$$\beta_{c1} \sim \text{Normal}(0, 0.1)$$

For each of four chains, there was a tuning phase of 2,000 iterations with a target acceptance rate of 0.9, followed by 10,000 samples. Convergence was assessed using bulk effective sample size and rank-normalized split potential scale reduction statistic $\hat{R}$ (Vehtari et al., 2021). We assessed the fit of the model using posterior predictive sampling of responses and response times. Finally, we calculated the 94% high-density interval for each of the group-level mean parameters to determine whether they were different from zero, indicating a relationship between a given neural signal and either the variable-speed or fixed-speed drift rate.

### Supplementary Results

#### *Age Group Differences in Performance and Angular Gyrus Activation*

Our main analyses examined developmental differences in behavioral performance and angular gyrus activation by treating age as a continuous variable. This approach allowed us to test whether performance and the relationship between neural activation and behavior changed systematically with increasing age. However, because our sample included children through age 12 years, but not older adolescents, the continuous-age approach assumes a linear developmental trajectory across an age range for which we do not have observations. To address this limitation, we conducted supplemental analyses treating age as a categorical variable. We re-ran the best-fitting model reported within the main text, replacing age in years with age group (children versus adults). Categorical variable effects were examined using deviation coding, such that comparisons were made to the grand mean instead of a specified reference level. Analyses were again conducted in R (R Core Team, 2026) using the same packages reported in the main text. As described below, the categorical analyses yielded a similar pattern of results to those treating age continuously.

**Initial associations provide stable premises for inference across development.** Memory for initial associations was reassessed using the main effects model (accuracy ~ age group + test phase + (1 I participant)). This model yielded a conditional $R^2 = 0.46$ and marginal $R^2 = 0.09$. Significant effects of age group ($\chi^2(1) = 7.35$, $p = .01$) and test phase ($\chi^2(1) = 6.50$, $p = .01$) were observed. Performance was slightly lower in children compared to adults. There was also a small decline in performance from initial learning to the final test across participants. Despite these slight differences, both children and adults formed and maintained robust memories of initial pairs (children: initial learning phase *M* = 98.04%, SEM = 0.36%, final test phase *M* = 96.60%, SEM = 0.71%; adults: initial learning phase *M* = 99.58%, SEM = 0.17%; final test phase *M* = 99.06%, SEM = 0.65%).

**Inference shows a disproportionate developmental improvement relative to memory.** We next reassessed memory versus inference performance. Accuracy was first examined using the interaction model (accuracy ~ age group + test type + age group × test type + (1 I participant)). This model yielded a conditional $R^2 = 0.68$ and marginal $R^2 = 0.28$. A significant age group x text phase interaction was observed ($\chi^2(2) = 16.73$, $p < .01$; age group effect: $\chi^2(1) = 14.01$, $p < .01$; test phase effect: $\chi^2(2) = 68.81$, $p < .01$). We interrogated this interaction using marginal contrasts. Comparisons between children and adults showed the greatest age group difference in inference performance ($t(214) = 5.31$, $p < .01$), followed by overlapping memory test performance ($t(214) = 2.48$, $p = .01$), and then non-overlapping memory test performance ($t(214) = 1.64$, $p = .10$).

We then examined test type performance in adults and children separately. Adults' inference performance was lower than both their overlapping and non-overlapping memory test performances ($ts(214) \leq -2.19$, $ps \leq .03$), which did not differ from one another, $t(214) = 0.13$, $p = .90$. Children demonstrated a similar pattern, though their inference performance was even lower compared to their overlapping and non-overlapping memory test performances ($ts(214) \leq -8.63$, $ps < .01$), which also did not differ from one another, $t(214) = -1.39$, $p = .17$. Participants were therefore less accurate on inference than memory for directly learned pairs, and this difference was largest in children, indicating a disproportionate difficulty in deriving relationships across associations earlier in development.

Response times for correct trials were then examined using the main effects model (response time ~ age group + test type + (1 I participant)). This model yielded a conditional $R^2 = 0.85$ and marginal $R^2 = 0.58$. Significant effects of age group ($\chi^2(1) = 36.72$, $p < .01$) and test phase ($\chi^2(2) = 629.39$, $p < .01$) were observed. Adults were faster to respond than children overall. Marginal contrasts also showed significantly longer response times for inference compared to both overlapping and non-overlapping memory tests ($ts(216) \geq 21.28$, $ps < .01$), which did not differ from one another ($t(216) = 0.87$, $p = .38$). Responses were therefore faster in adults versus children across all conditions, but remained reliably slower for inference than memory.

**Angular gyrus engagement indexes increasingly efficient inference with age.** Angular gyrus activation was then reassessed using the interaction model (angular gyrus PE ~ age group + test type + age group × test type + (1 I participant)). This model yielded a marginal $R^2 = 0.10$. A significant age group x test phase interaction was observed ($\chi^2(1) = 9.78$, $p < .01$; main effect of age group: $\chi^2(1) = 3.69$, $p = .05$; main effect of test type: $\chi^2(1) = 0.94$, $p = .33$). We interrogated this interaction using marginal contrasts, comparing children and adults' performance. Results showed significantly greater recruitment of the angular gyrus during inference in adults versus children ($t(120) = 3.57$, $p < .01$). In contrast, the two age groups did not differ in their recruitment of the angular gyrus during memory ($t(120) = -0.85$, $p = .40$).

We next reassessed whether angular gyrus engagement during inference predicted faster correct inference using the interaction model (response time ~ age group + angular gyrus PE + age group × angular gyrus PE). This model explained substantial variance (adjusted $R^2 = 0.33$, $F(3,59) = 11.02$, $p < .01$). An age group x angular gyrus PE interaction was observed ($F(1) = 3.91$, $p = .05$; main effect of age: $F(1) = 20.77$, $p < .01$; main effect of angular gyrus PE: $F(1) = 0.19$, $p = .66$). In adults, greater angular gyrus engagement tended to predict faster correct inference. The opposite was observed in children, for whom angular gyrus engagement tended to predict slower responses. This pattern is consistent with a role of angular gyrus in efficient, direct inference in adults but not children.

#### *Exploratory Response Time Model Results for Additional Neural Regions*

To assess the specificity of the primary findings, we conducted exploratory analyses examining whether additional regions previously implicated in adult inference and memory-based reasoning (Morton et al., 2020; Zeithamova & Preston, 2010) tracked the computational processes identified by the model. These regions included ventrolateral prefrontal cortex (VLPFC), intraparietal sulcus/superior parietal lobule (IPS/SPL), precuneus, and parahippocampal cortex (PHC).

Activation in VLPFC and IPS/SPL predicted slower response times associated with the variable-speed process across age groups (**Fig. S1**). These regions were not reliably related to the speed of the fixed-speed process on average. However, both showed age-related changes in this relationship. In adults, greater activation in VLPFC and IPS/SPL was associated with slower evidence accumulation for the fixed-speed process. This pattern suggests that these regions may support control-related processes engaged when retrieval is effortful. Such processes may be recruited to facilitate iterative retrieval across development and, in adults, may also be engaged when more efficient, direct retrieval routes are unavailable or fail.

A distinct pattern emerged in precuneus and PHC. Activation in these regions was associated with faster operation of the variable-speed process in adults, although this effect was only reliable in precuneus (**Fig. S1**). In contrast, activation in both regions was associated with slower and less accurate decisions linked to the fixed-speed process. This pattern is consistent with the possibility that precuneus and PHC contribute to representing contextual information that supports retrieval of specific event details, thereby facilitating iterative retrieval. At the same time, increased engagement of these regions may reflect conditions under which direct, integrated representations are not successfully accessed, leading to slower and less accurate performance on the fixed-speed process.

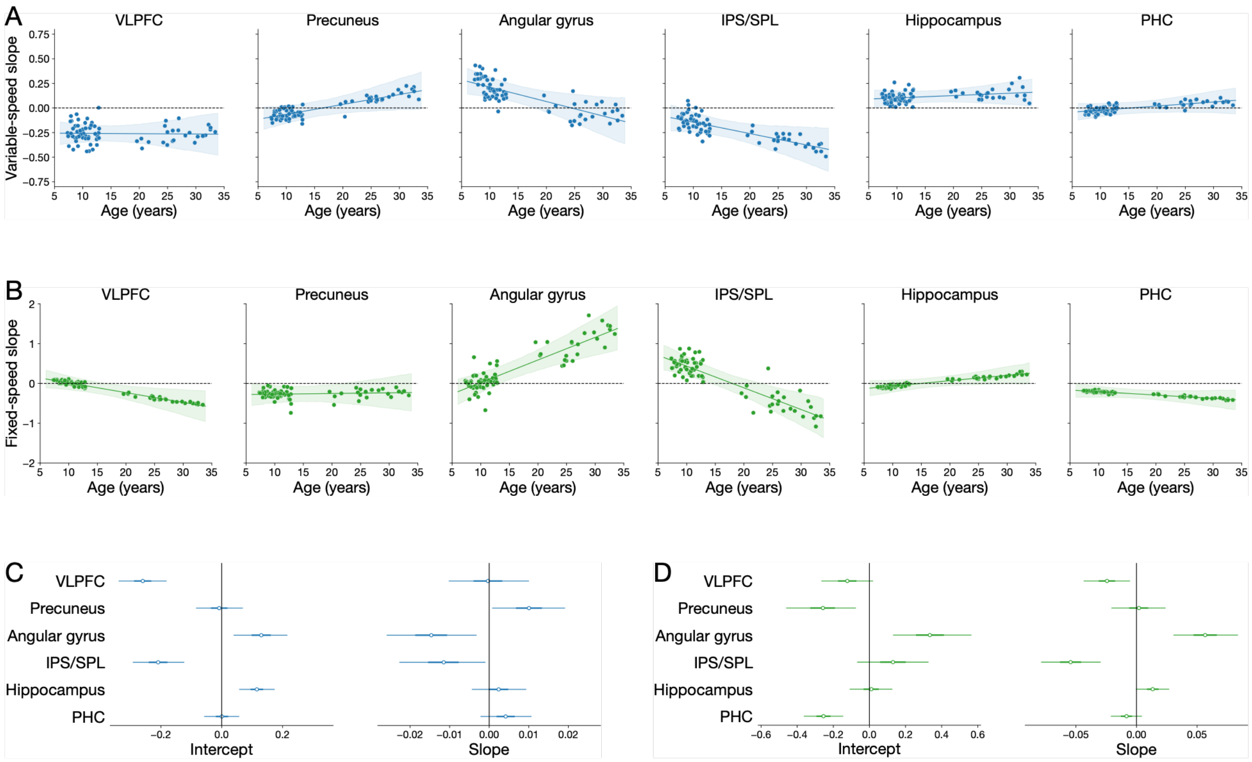


**Figure S1.** Neural modulation of accumulator drift rates across development. Results are shown for the hippocampus and angular gyrus (identical to those in the main manuscript), as well as for regions examined in exploratory analyses. **(A)** Subject-specific neural slope parameters relating trial-by-trial neural activation to drift rate for the variable-speed accumulator ($v_1$). **(B)** Subject-specific neural slope parameters for the fixed-speed accumulator ($v_2$). **(C)** Age-related effects (regression coefficients) on neural slope parameters for the variable-speed accumulator. **(D)** Age-related effects on neural slope parameters for the fixed-speed accumulator. Positive values indicate that greater neural activation is associated with faster evidence accumulation, whereas negative values indicate slower accumulation.

#### ***Model Convergence, Parameter Estimates, and Fit***

We first assessed convergence of the Bayesian sampling procedure. There were no divergences during sampling. For all model parameters, the rank-normalized split potential scale reduction statistic was low ($\hat{R} < 1.0064$), and effective sample sizes were at least 1007, indicating successful convergence and sufficient sampling for stable parameter estimation.

Posterior estimates of group-level parameters are reported in **Table S3**. Subject-specific estimates of decision threshold and mean accumulator drift rates are shown in **Fig. S2**. Age-related effects on neural slope parameters are reported in **Table S4** and visualized in **Fig. S1**, and corresponding standard deviations are reported in **Table S5**.

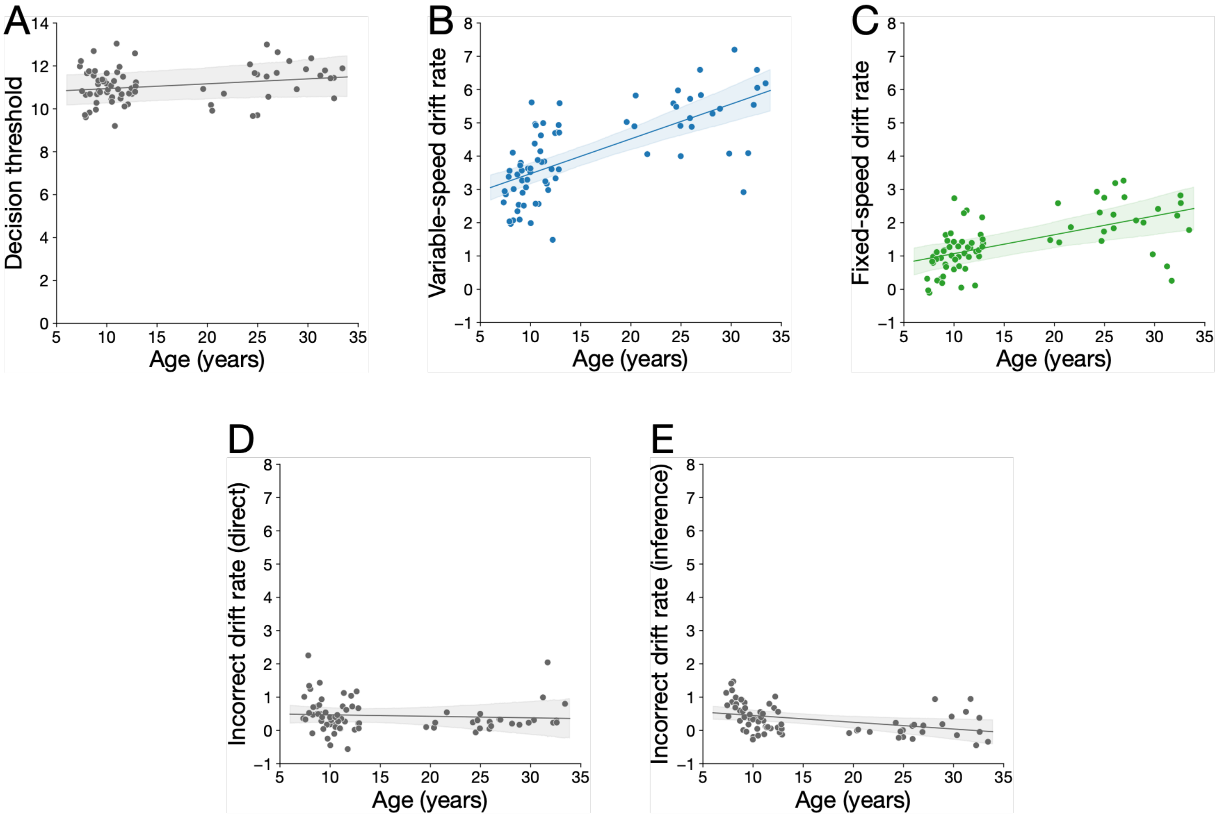


**Figure S2.** Subject-specific parameter estimates as a function of age. **(A)** Decision threshold parameter ($b$). **(B)** Variable-speed drift rate ($v_1$). **(C)** Fixed-speed drift rate ($v_2$). **(D)** Drift rate for incorrect responses on memory tests ($v_3$). **(E)** Drift rate for incorrect responses on inference tests ($v_4$). Each point reflects a subject-specific posterior estimate.

To evaluate model fit, we used posterior predictive sampling, in which parameters drawn from the posterior distribution were used to simulate responses and response times. We first assessed mean response accuracy, averaged across posterior samples, and found that the model captured individual differences in accuracy across participants (**Fig. S3**).

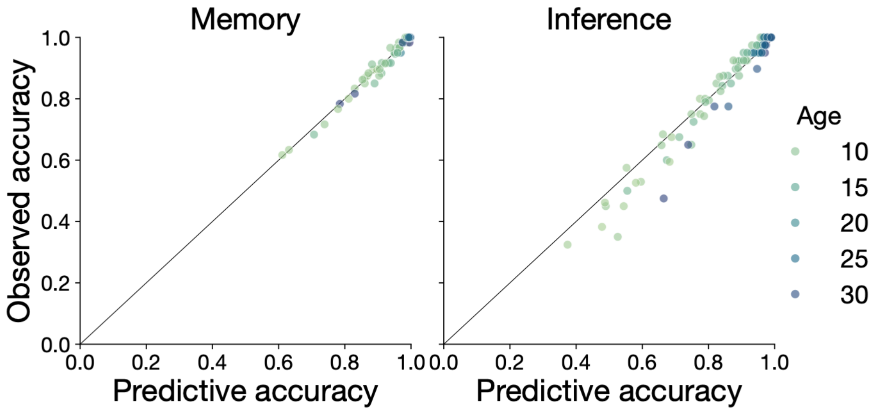


**Figure S3.** Observed and simulated accuracy for memory and inference tests. Simulated responses were generated using posterior predictive sampling from the fitted model. Points reflect observed and predicted accuracy for individual participants.

We next examined predicted response times for correct and incorrect responses on memory and inference trials. In rare cases, posterior samples produced mean response times exceeding the response deadline (8 s); these samples were excluded. Overall, simulated response times closely matched observed individual differences across conditions (**Fig. S4**).

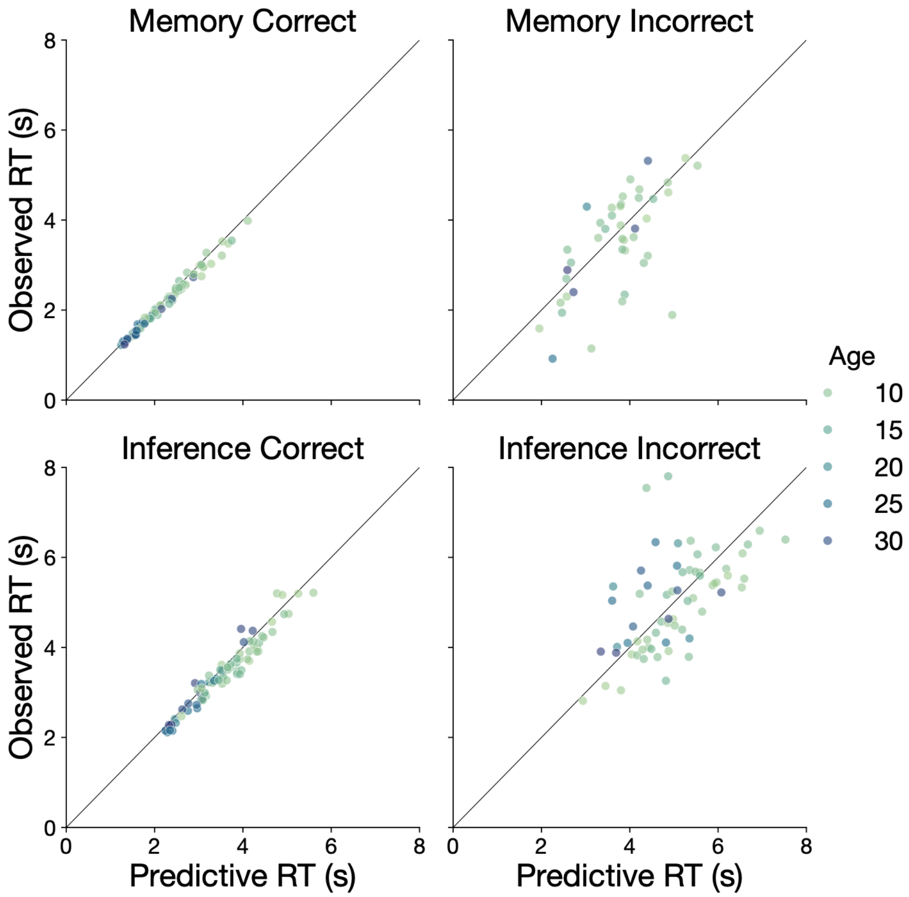


**Figure S4.** Observed and simulated response times for correct and incorrect responses on memory and inference tests. Simulated responses were generated using posterior predictive sampling from the fitted model. Points reflect observed and predicted response times for individual participants.

To assess fit at the trial level, we compared observed response times on individual trials to the mean predicted response time for those trials, averaged across posterior samples. As above, samples producing implausibly slow responses were excluded. The model captured trial-level variability in response time (**Fig. S5**).

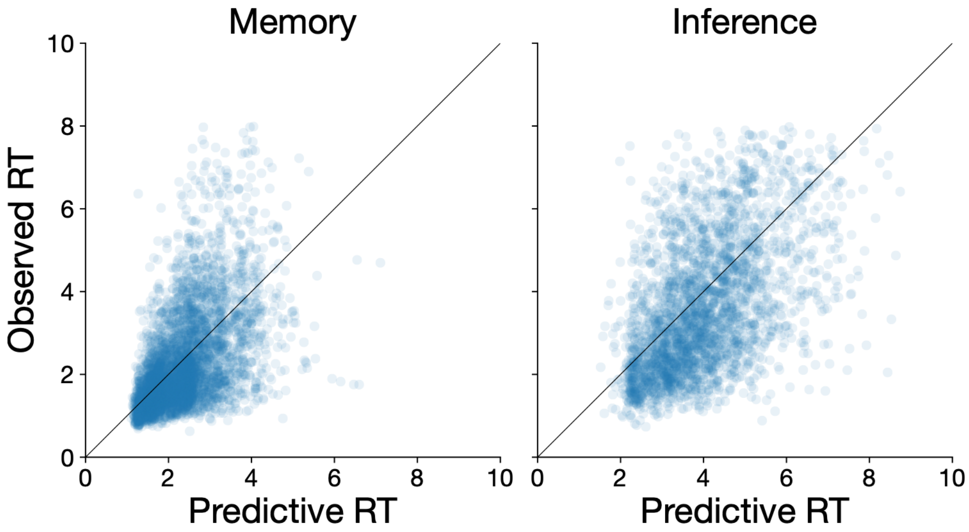


**Figure S5.** Observed and simulated response times on individual trials. Simulated responses were generated using posterior predictive sampling from the fitted model. Points reflect observed and predicted response times at the trial level.

Finally, we compared observed and simulated response time distributions across age groups. For memory tests, response time distributions were well captured by posterior predictive samples across all age groups (**Fig. S6**). For inference tests, distributions were also generally well fit, although incorrect responses in adults were somewhat faster in the model than in the observed data (**Fig. S7**).

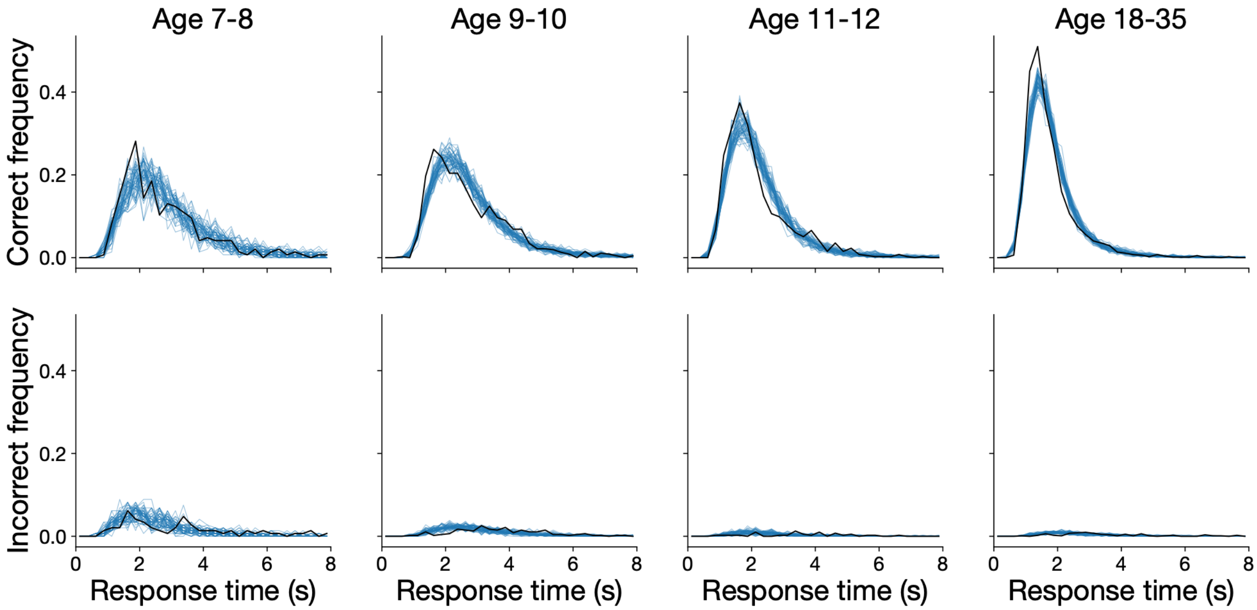


**Figure S6.** Observed and simulated response time distributions for memory tests across age groups. Response time distributions are shown as relative frequencies for each age group. Simulated distributions were generated using posterior predictive sampling from the fitted model. The height of each distribution is scaled by the fraction of trials with that response (correct or incorrect). Black lines indicate observed response times; blue lines indicate 50 posterior predictive samples.

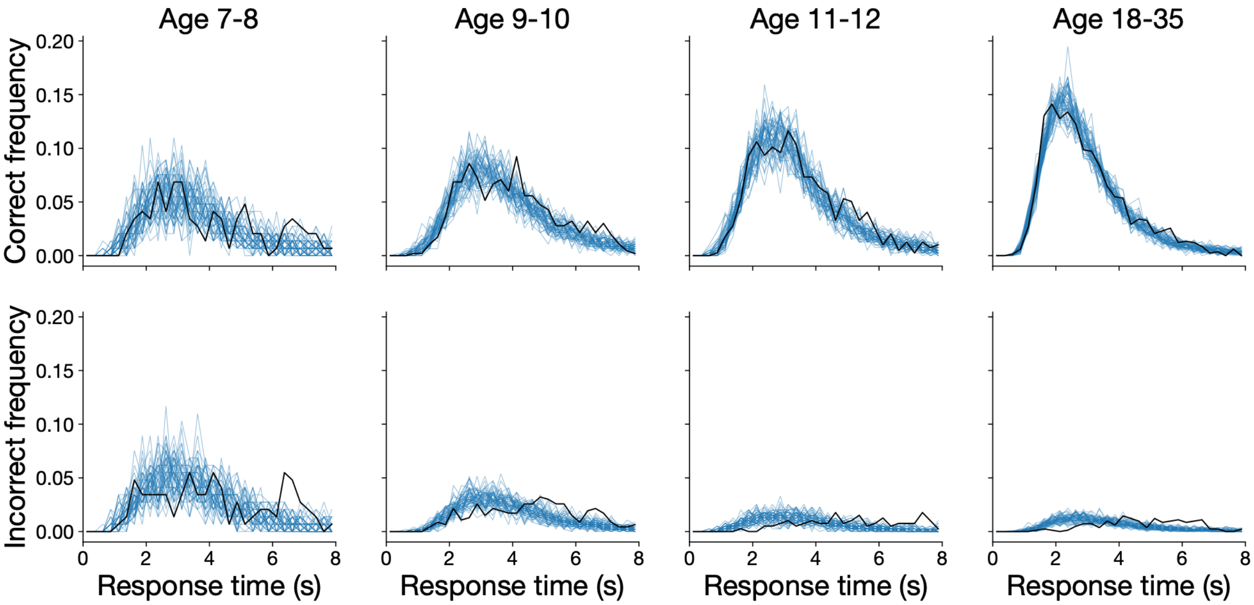


**Figure S7**. Observed and simulated response time distributions for inference tests across age groups. Response time distributions are shown as relative frequencies for each age group. Simulated distributions were generated using posterior predictive sampling from the fitted model. The height of each distribution is scaled by the fraction of trials with that response (correct or incorrect). Black lines indicate observed response times; blue lines indicate 50 posterior predictive samples.

**Table S1.** Statistically Significant fMRI Clusters (anatomic labels from Harvard-Oxford Atlas).

| Peak MNI coordinates | | | Max intensity | Cluster size | Anatomical region at cluster peak |
|---|---|---|---|---|---|
| x | y | z | value (z-stat) | (voxels) | |
| **Main effect of inference > memory within hippocampus[2]** | | | | | |
| 18 | -9 | -26 | 2.675 | 360 | 73% Parahippocampal Gyrus; 25% Right Hippocampus |
| -11 | -9 | -23 | 2.425 | 304 | 39% Parahippocampal Gyrus; 16% Left Hippocampus |
| 33 | -29 | -14 | 3.061 | 162 | 50% Right Hippocampus; 15% Parahippocampal Gyrus |
| -21 | -38 | -5 | 2.414 | 131 | 43% Left Hippocampus |
| 16 | -34 | -6 | 2.450 | 119 | 40% Parahippocampal Gyrus; 10% Lingual Gyrus; 10% Right Hippocampus |
| **Main effect of inference > memory across whole brain[1]** | | | | | |
| 33 | -9 | -41 | 2.363 | 117025 | 81% Temporal Fusiform Cortex; 7% Inferior Temporal Gyrus |
| -47 | 31 | 14 | 2.340 | 17986 | 52% Inferior Frontal Gyrus; 8% Middle Frontal Gyrus; 7% Frontal Pole |
| 12 | 21 | 29 | 2.763 | 8112 | 32% Cingulate Gyrus; 14% Paracingulate Gyrus |
| 55 | 32 | 14 | 2.513 | 5640 | 49% Inferior Frontal Gyruss; 17% Frontal Pole; 6% Middle Frontal Gyrus |
| -1 | 38 | -24 | 2.416 | 4166 | 96% Frontal Medial Cortex |
| 30 | 23 | -10 | 2.375 | 2973 | 64% Frontal Orbital Cortex; 11% Insular Cortex |
| -40 | 46 | -17 | 2.437 | 2632 | 56% Frontal Pole |
| -28 | 20 | -10 | 2.487 | 2167 | 49% Frontal Orbital Cortex; 25% Insular Cortex |
| -56 | -50 | -18 | 2.319 | 1879 | 67% Inferior Temporal Gyrus; 5% Middle Temporal Gyrus |
| -15 | -3 | -8 | 2.676 | 1812 | 51% Left Pallidum |
| 29 | 9 | -5 | 2.670 | 1215 | 58% Right Putamen |
| **Positive effect of age on inference > memory across whole brain[1]** | | | | | |
| 63 | -21 | -31 | 2.311 | 31002 | 39% Inferior Temporal Gyrus; 5% Middle Temporal Gyrus |
| -55 | -34 | 30 | 2.478 | 19735 | 32% Supramarginal Gyrus; 17% Parietal Operculum Cortex |
| -23 | -90 | -21 | 3.163 | 13285 | 18% Occipital Fusiform Gyrus; 10% Lateral Occipital Cortex; 6% Occipital Pole |
| -27 | 46 | -13 | 2.32 | 4520 | 88% Frontal Pole |
| -41 | -41 | -28 | 2.334 | 2510 | 37% Temporal Fusiform Cortex; 18% Inferior Temporal Gyrus; 10% Temporal Occipital Fusiform Cortex |
| 36 | -5 | 43 | 3.061 | 2124 | 21% Precentral Gyrus; 10% Middle Frontal Gyrus |
| 48 | 36 | 17 | 2.376 | 1822 | 40% Frontal Pole; 19% Middle Frontal Gyrus; 15% Inferior Frontal Gyrus |
| -55 | 6 | 10 | 2.381 | 1784 | 69% Precentral Gyrus; 13% Inferior Frontal Gyrus |
| -62 | -29 | -28 | 2.586 | 1212 | 46% Inferior Temporal Gyrus |
| 69 | -36 | -3 | 2.335 | 1037 | 76% Middle Temporal Gyrus |

[1]Whole-brain cluster-corrected for multiple comparisons, $p < .01$; $a < .05$.

[2]Small-volume cluster corrected for multiple comparisons, $p < .01$; $a < .05$.

**Table S2**. Posterior estimates of slope parameters for children and adults, and high-density intervals for the difference between adults and children.

| Parameter | Child | Adult | Adult difference 94% HDI |
|---|---|---|---|
| $v_{1,\mathrm{vlpfc}}$ | -0.254702 | -0.270476 | [-0.1710, 0.1410] |
| $v_{2,\mathrm{vlpfc}}$ | 0.013390 | -0.406886 | [-0.7355, -0.1036] |
| $v_{1,\mathrm{prec}}$ | -0.063704 | 0.107082 | [0.0267, 0.3209] |
| $v_{2,\mathrm{prec}}$ | -0.270099 | -0.240622 | [-0.3306, 0.3972] |
| $v_{1,\mathrm{ang}}$ | 0.211923 | -0.040233 | [-0.4279, -0.0772] |
| $v_{2,\mathrm{ang}}$ | 0.021910 | 0.997590 | [0.5664, 1.3886] |
| $v_{1,\mathrm{lpc}}$ | -0.148107 | -0.336654 | [-0.3579, -0.0158] |
| $v_{2,\mathrm{lpc}}$ | 0.439416 | -0.513212 | [-1.3265, -0.5773] |
| $v_{1,\mathrm{hpc}}$ | 0.102413 | 0.143586 | [-0.0650, 0.1464] |
| $v_{2,\mathrm{hpc}}$ | -0.064946 | 0.163662 | [0.0038, 0.4511] |
| $v_{1,\mathrm{phc}}$ | -0.021658 | 0.048181 | [-0.0327, 0.1746] |
| $v_{2,\mathrm{phc}}$ | -0.210040 | -0.348713 | [-0.3542, 0.0713] |

**Table S3.** Posterior estimates of group-level parameters.

| Parameter | Mean | 94% HDI |
|---|---|---|
| $\tau$ | 0.006845 | [0.0000, 0.0192] |
| $A$ | 5.904910 | [5.3307, 6.5190] |
| $b_{\beta 0}$ | 2.403807 | [2.3416, 2.4670] |
| $b_{\beta 1}$ | 0.002041 | [-0.0006, 0.0047] |
| $\sigma_b$ | 0.084487 | [0.0671, 0.1022] |
| $v_{1,\beta 0}$ | 4.057317 | [3.7731, 4.3402] |
| $v_{1,\beta 1}$ | 0.104611 | [0.0764, 0.1338] |
| $\sigma_1$ | 1.008555 | [0.8277, 1.1954] |
| $v_{2,\beta 0}$ | 1.387986 | [1.0301, 1.7336] |
| $v_{2,\beta 1}$ | 0.056631 | [0.0296, 0.0832] |
| $\sigma_2$ | 0.810247 | [0.6098, 1.0167] |
| $r$ | 0.472482 | [0.4357, 0.5060] |
| $v_{3,\beta 0}$ | 0.443789 | [0.2226, 0.6577] |
| $v_{3,\beta 1}$ | -0.004621 | [-0.0305, 0.0214] |
| $\sigma_3$ | 0.641296 | [0.4744, 0.8219] |
| $v_{4,\beta 0}$ | 0.335089 | [0.1881, 0.4862] |
| $v_{4,\beta 1}$ | -0.020396 | [-0.0375, -0.0040] |
| $\sigma_4$ | 0.488063 | [0.3793, 0.6017] |

**Table S4.** Posterior estimates of neural signal slope age coefficients.

| Parameter | Mean | 94% HDI |
|---|---|---|
| $v_{1,\mathrm{vlpfc},\beta 0}$ | -0.259739 | [-0.3387, -0.1809] |
| $v_{1,\mathrm{vlpfc},\beta 1}$ | -0.000344 | [-0.0102, 0.0100] |
| $v_{1,\mathrm{prec},\beta 0}$ | -0.008307 | [-0.0843, 0.0696] |
| $v_{1,\mathrm{prec},\beta 1}$ | 0.010060 | [0.0008, 0.0192] |
| $v_{1,\mathrm{ang},\beta 0}$ | 0.130047 | [0.0391, 0.2161] |
| $v_{1,\mathrm{ang},\beta 1}$ | -0.014661 | [-0.0259, -0.0032] |
| $v_{1,\mathrm{lpc},\beta 0}$ | -0.209147 | [-0.2922, -0.1231] |
| $v_{1,\mathrm{lpc},\beta 1}$ | -0.011573 | [-0.0227, -0.0010] |
| $v_{1,\mathrm{hpc},\beta 0}$ | 0.115601 | [0.0574, 0.1741] |
| $v_{1,\mathrm{hpc},\beta 1}$ | 0.002375 | [-0.0044, 0.0094] |
| $v_{1,\mathrm{phc},\beta 0}$ | 0.001115 | [-0.0576, 0.0575] |
| $v_{1,\mathrm{phc},\beta 1}$ | 0.004155 | [-0.0022, 0.0107] |
| $v_{2,\mathrm{vlpfc},\beta 0}$ | -0.122821 | [-0.2663, 0.0200] |
| $v_{2,\mathrm{vlpfc},\beta 1}$ | -0.024531 | [-0.0436, -0.0053] |
| $v_{2,\mathrm{prec},\beta 0}$ | -0.260026 | [-0.4613, -0.0747] |
| $v_{2,\mathrm{prec},\beta 1}$ | 0.001754 | [-0.0207, 0.0240] |
| $v_{2,\mathrm{ang},\beta 0}$ | 0.337461 | [0.1305, 0.5649] |
| $v_{2,\mathrm{ang},\beta 1}$ | 0.056872 | [0.0305, 0.0837] |
| $v_{2,\mathrm{lpc},\beta 0}$ | 0.130056 | [-0.0681, 0.3275] |
| $v_{2,\mathrm{lpc},\beta 1}$ | -0.054498 | [-0.0787, -0.0296] |
| $v_{2,\mathrm{hpc},\beta 0}$ | 0.009215 | [-0.1093, 0.1271] |
| $v_{2,\mathrm{hpc},\beta 1}$ | 0.013442 | [-0.0006, 0.0270] |
| $v_{2,\mathrm{phc},\beta 0}$ | -0.254859 | [-0.3635, -0.1453] |
| $v_{2,\mathrm{phc},\beta 1}$ | -0.008355 | [-0.0212, 0.0045] |

**Table S5.** Posterior estimates of neural slope standard deviations.

| Parameter | Mean | 94% HDI |
|---|---|---|
| $\sigma_{1,\mathrm{vlpfc}}$ | 0.163554 | [0.0733, 0.2464] |
| $\sigma_{1,\mathrm{prec}}$ | 0.105995 | [0.0000, 0.1903] |
| $\sigma_{1,\mathrm{ang}}$ | 0.173588 | [0.0821, 0.2652] |
| $\sigma_{1,\mathrm{lpc}}$ | 0.142732 | [0.0329, 0.2380] |
| $\sigma_{1,\mathrm{hpc}}$ | 0.109786 | [0.0171, 0.1816] |
| $\sigma_{1,\mathrm{phc}}$ | 0.074766 | [0.0000, 0.1395] |
| $\sigma_{2,\mathrm{vlpfc}}$ | 0.128825 | [0.0000, 0.2732] |
| $\sigma_{2,\mathrm{prec}}$ | 0.256637 | [0.0017, 0.4301] |
| $\sigma_{2,\mathrm{ang}}$ | 0.429819 | [0.2121, 0.6459] |
| $\sigma_{2,\mathrm{lpc}}$ | 0.402902 | [0.2236, 0.5942] |
| $\sigma_{2,\mathrm{hpc}}$ | 0.102213 | [0.0000, 0.2267] |
| $\sigma_{2,\mathrm{phc}}$ | 0.079204 | [0.0000, 0.1800] |
